# Molecular Imaging with Fibroblast Activation Protein Tracers depicts Inflammatory Joint Damage and its Transition to Resolution of Inflammation

**DOI:** 10.1101/2023.01.17.524425

**Authors:** Simon Rauber, Hashem Mohammadian, Christian Schmidkonz, Armin Atzinger, Alina Soare, Simone Maschauer, Christoph Treutlein, Mario Angeli, Maria Gabriella Raimondo, Cong Xu, Kai-Ting Yang, Le Lu, Hannah Labinsky, Eleni Kampylafka, Johannes Knitza, Hans Maric, Jörg H.W. Distler, Tobias Bäuerle, Torsten Kuwert, Olaf Prante, Juan Cañete, Georg Schett, Andreas Ramming

## Abstract

Joint fibroblasts play an important role in the transition from joint inflammation to irreversible joint damage. There is no established clinical method to measure fibroblast activation during inflammation and their phenotypic dynamics upon therapy to date. Here we show that upon treatment with IL-17A/TNF-blocking antibodies fibroblasts change their phenotype from a destructive IL-6^+^/MMP3^+^THY1^+^ to a CD200^+^DKK3^+^ subtype, actively inducing resolution of inflammation. This phenotypic switch can be visualized due to so far unexplored different capacities of fibroblast subtypes with regard to receptor internalization of small molecular tracers with high affinity to FAP. Although FAP expression levels are comparable between fibroblast subtypes in the joint, FAP internalisation rate correlates with the destructive potential of fibroblasts and resolving fibroblasts have a lower FAP internalisation rate, providing a valuable imaging tool to visualize the transition from joint damage to resolution of inflammation.

## INTRODUCTION

Tissue damage in joints directly leads to functional impairment and disability because of very limited repair capacities of the joint cartilage and the adjacent bone^1,2^. The most common trigger of irreversible cartilage and bone destruction are chronic inflammatory processes in the joints. Timely limitation of inflammation seems to be essential to prevent joint damage and is therefore main target of immunosuppressive therapies ^3,4^. Despite significant progress in the development of targeted biologic therapies, more than 40% of patients still record gradual loss of joint function, mostly because of inflammatory flares and persistent disease activity during therapy. Conversely, patients of all common entities including rheumatoid arthritis, mixed connective tissue diseases as well as spondyloarthropathies also present non-deforming courses of joint inflammation. While in the latter patients, current anti-inflammatory treatment is entirely sufficient, patients with persistent inflammation and relapses require therapy adjustment and escalation until mesenchymal activation is quieted.

To date there is no diagnostic tool available to decide whether inflammation is damaging the joint. Regular magnetic resonance imaging (MRI) or ultrasound examinations are normally carried out to detect silent damage over time. This approach results into a delay of intervention as treatment escalation can only be initiated after the damage has already occurred.

It was shown that the depletion of activated synovial fibroblasts prevented joint damage despite presence of a residual inflammation ^5^. Fibroblasts therefore appear to play a key role in the transition from inflammation to joint damage, and measuring fibroblast activity in the joint might be a valuable tool to improve treatment outcomes^6^.

New small molecular tracers expand the spectrum of measurable cell activities ^7,8^. Quinolone-based tracers that act as highly specific ligands of human and mouse fibroblast activation protein (FAP), so called FAPIs, showed high affinities for activated fibroblasts in various studies ^8^. Using Positron emission tomography (PET) imaging, ^68^Ga-loaded FAPI-04 has been positively validated to dissect inflammatory from fibroblast activity in human disease ^9^. This tracer also binds to synovial fibroblasts with high affinity in chronic inflammatory joint diseases ^10^.

Here we show that FAPI-04 tracer uptake in inflamed joints is associated with joint damage and modulated upon targeted biologic treatment. However, the respective FAPI-04 uptake does not simply reflect the total fibroblast activity level in the respective joint, but rather visualizes the current presence of functionally divergent fibroblast subtypes. Whereas inflammatory, destructive fibroblasts internalize FAPI-04 at high rates, resolution of inflammation is associated with a phenotypic transition into CD200/DKK3^+^ fibroblasts, internalizing FAPI-04 at significantly lower rates. This so far unknown mesenchymal cell fate protects from inflammation via CD200/CD200R and activates innate lymphoid cells ^11^, which are known to be essential cellular components for the activation of regulatory T cells and pro-resolving eosinophilic granulocytes in the joint ^12^. The correlation between FAPI-04 uptake and presence of functionally distinct fibroblast subtypes can be used not only to visualize active joint damage but might also improve drug monitoring and silent joint destruction under DMARD therapies in particular in patients with disease flares and persistent inflammatory disease activity.

## RESULTS

### FAPI-04 uptake in patients with immune-mediated inflammatory diseases

Fibroblasts are important mediators to induce joint damage, however measuring mesenchymal tissue activation in arthritis patients is challenging but may represent a possibility to detect regions at risk for structural damage. PET imaging with FAPI tracers has been shown to be a valuable tool to detect mesenchymal activity independent of inflammatory processes ^9^. In immune-mediated inflammatory diseases (IMIDs) we observed ^68^Ga-FAPI-04 accumulation at synovial and enthesial sites compared to non-arthritic controls **(Fig. 1a,b)**. Signal intensities were similar in peripheral and axial anatomical regions and did not differ with regard to the affected tissue **(Extended Data Fig. 1)**. ^68^Ga-FAPI-04 uptake concurred with signs of inflammation as measured by simultaneous magnetic resonance imaging (MRI) and correlated with several composite scores measuring clinical disease activity **(Fig. 1c,d)**. Therefore, inflamed regions were further stratified based on their respective signs of structural damage. Lesions with erosive defects, inflamed regions demonstrating osteoproliferative changes of the tissue, as well as inflammation sites without any signs of inceptive damage were included into the analysis **(Fig. 1d)**. While ^68^Ga-FAPI-04 uptake was absent in defect-free inflamed areas, a significant uptake was detected in both erosive as well as osteoproliferative lesions. Anti-inflammatory treatment with tumor necrosis factor alpha (TNF) inhibitors or interleukin (IL)-17A blocking antibodies resulted into decline of FAPI-04 accumulation **(Fig. 1e)**. In a sex and age matched comparative analysis, the reduction of FAPI-04 was significantly stronger in the anti-IL-17A treated compared to the anti-TNF treated group.

**Figure 1:**
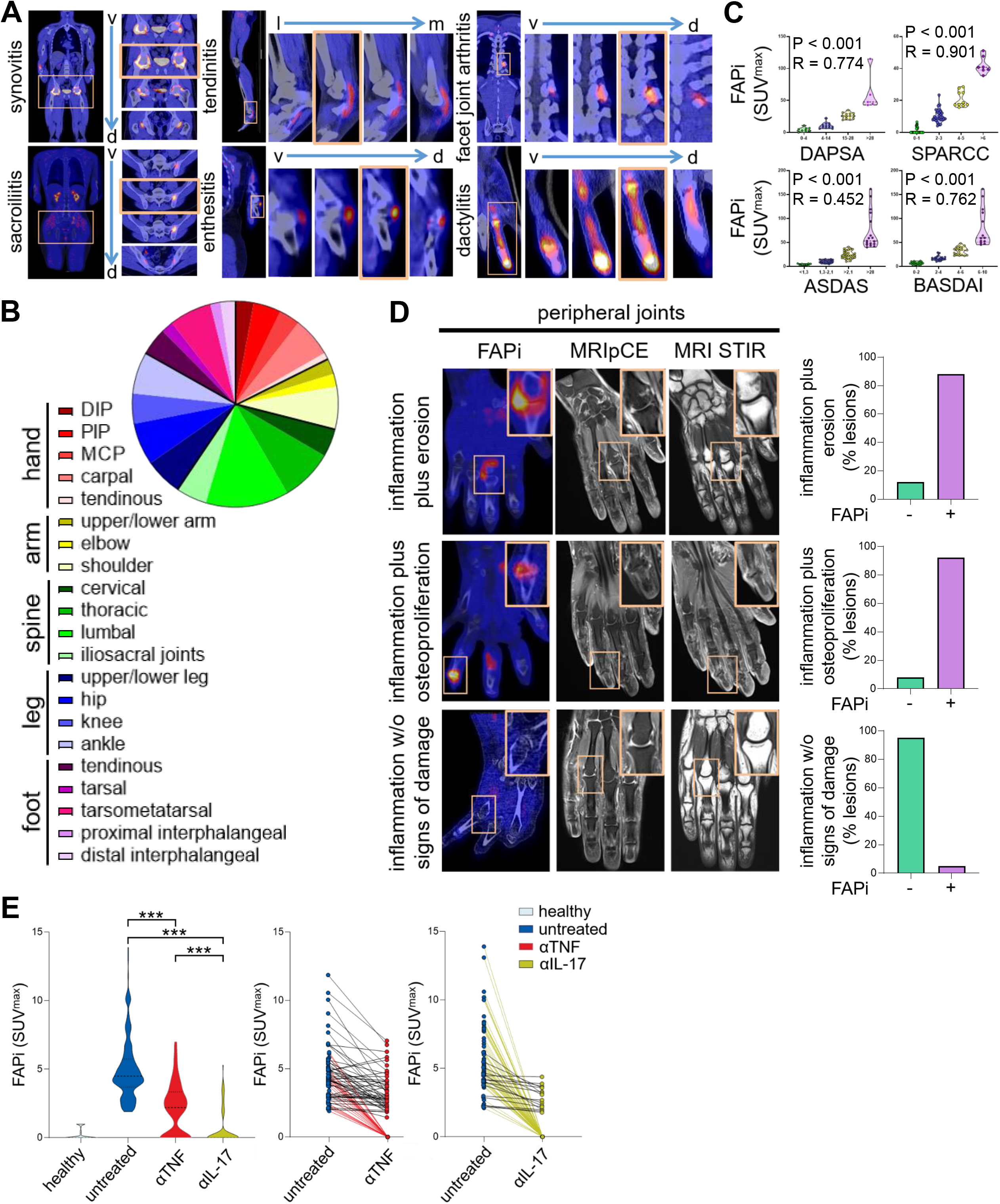
FAPI-04 uptake in patients with immune-mediated inflammatory diseases. **(a)** Representative PET/CT scans with ^68^Ga-FAPI-04 tracers and their accumulation sites in patients with immune-mediated inflammatory diseases (IMIDs) including rheumatoid arthritis (N = 20), psoriatic arthritis (N = 85), axial spondyloarthritis (N = 28). Four consecutive sectional planes of higher magnifications of the marked areas are shown from ventral (v) to dorsal (d) or lateral (l) to medial (m). **(b)** Body distribution of ^68^Ga-FAPI-04 positive lesions of the whole cohort of IMID patients. **(c)** Pearson correlation analysis of ^68^Ga-FAPI-04 uptake (maximum standardized uptake value; SUV^max^) and composite scores of clinical/radiological disease activities of the respective IMIDs; disease activity in psoriatic arthritis (DAPSA); spondyloarthritis research consortium of Canada scoring (SPARCC); ankylosing spondylitis disease activity (ASDAS); Bath ankylosing spondylitis disease activity (BASDAI); **(d)** Comparative analysis of ^68^Ga-FAPI-04 PET/CT and MRI scans from the same patients (N = 30); lesions that showed signs of inflammation by MRI according to radiologist’s assessement, were separated into 3 groups: (i) inflammation plus signs of erosive joint damage; (ii) inflammation plus signs of osteoproliferative joint damage; (iii) inflammation without any sign of joint damage as assessed by MRI; lesions showing ^68^Ga-FAPI-04 were calculated. **(e)** ^68^Ga-FAPI-04 tracer uptake before and after treatment with TNF (N = 15) or IL-17A (N = 19). Non-arthritic individuals who underwent PET/CT served as controls (N = 15). ***p<0.001 determined by one-way ANOVA with Tukey’s multiple comparison post hoc test.

### FAP uptake correlates with joint damage

To examine the physiologic relevance of FAPI-04 uptake in joints, different mouse models of arthritis were screened by PET imaging for their respective ^68^Ga-FAPI-04 uptake and histologically analyzed for signs of inflammation and damage **(Fig. 2a-f)**. Wildtype animals showed no FAPI-04 uptake and a preserved architecture of joints. Human TNF transgenic (hTNFtg)197 mice (8-10 weeks old) and mice challenged with *I123* minicircles (mc) for 9 days prior to imaging accumulated FAPI-04 at moderate levels in ankle joints which was associated with a mild inflammation and sparse defects of bone architecture adjacent to the joint. Distinct levels of FAPI-04 accumulation were observed in mice overexpressing hTNF and mIL-23 correlating with the grade of inflammation and damage of the respective ankle joint. There was no difference of FAPI-04 uptake in erosive as compared to osteoproliferative lesions **(Fig. 2f)**. Similar to humans, treatment with antibodies against TNF and IL-17a significantly reduced articular FAPI-04 uptake **(Fig. 2g-i)**. Thereby, the superiority of IL-17a blockers in reducing FAPI-04 uptake was demonstrated across species and correlated with a reduction of joint damage.

**Figure 2:**
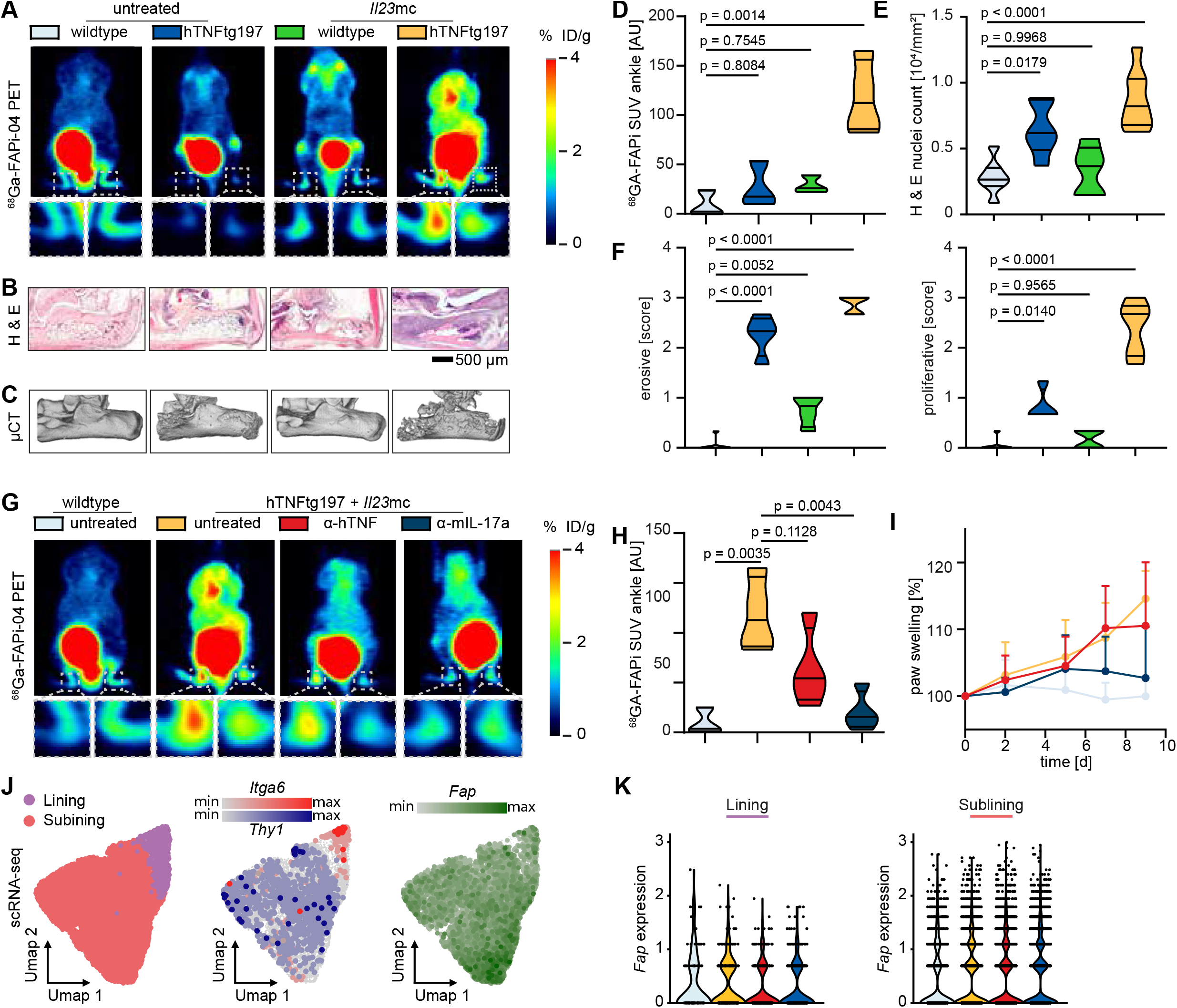
^68^Ga-FAPi-04 PET imaging in mice correlates with joint damage. **(a-c)** Representative images of **(a)** ^68^Ga-FAPI-04 PET imaging, **(b)** hematoxylin & eosin (H & E) stained ankle sections, **(c)** μCT images of wildtype animals and hTNFtg197 treated with Il23mc for 9 days prior to imaging; **(d)** Specific uptake of ^68^Ga-FAPI-04 tracer in ankle joints of animals corresponding to (a); median + quartiles; N=3 (untreated wildtype), N=3 (untreated hTNFtg197), N=3 (I123mc wildtype), N=4 (*I123*mc hTNFtg197). **(e)** Quantification of the number of nuclei / mm^2^ in ankle sections of animals corresponding to (a); median + quartiles; N=7 (untreated wildtype), N=6 (untreated hTNFtg197), N=5 (*I123*mc wildtype), N=8 (*I123*mc hTNFtg197). **(f)** μCT based erosion and proliferation scores for animals corresponding to (a); median + quartiles; N=6 (untreated wildtype), N=4 (untreated hTNFtg197), N=4 (*I123*mc wildtype), N=5 (*I123*mc hTNFtg197). **(g)** Representative images of ^68^Ga-FAPi-04 PET imaging of wildtype animals and hTNFtg187 treated with Il23mc for 9 days receiving either no treatment, anti-hTNF or anti-mIL-17a treatment. **(h)** Specific uptake of ^68^Ga-FAPi-04 tracer in ankle joints for animals corresponding to (g); median + quartiles; N=3 (untreated wildtype), N=4 (hTNFtg197 + *I123*mc untreated), N=4 (hTNFtg197 + *I123*mc +anti-hTNF), N=4 (hTNFtg197 + *I123*mc + anti-mIL-17a). **(i)** Clinical score (paw swelling) over time of animals corresponding to (g); mean + SD; N = 12 for all groups. **(j)** U-maps of a scRNA-seq dataset generated from sorted fibroblasts of wildtype animals and hTNFtg197 treated with Il23mc for 9 days receiving either no treatment, anti-hTNF or anti-mIL-17a treatment. U-maps show separation of lining and sublining fibroblasts, the expression of corresponding marker genes (*Itga6* and *Thy1,* respectively) and the expression of *Fap.* N=3 per condition. **(k)** *Fap* expression of lining and sublining fibroblasts between conditions.

### FAP expression across all fibroblast subtypes independent of anti-inflammatory treatment

To further investigate the underlying molecular mechanisms of decreasing FAPI uptake upon treatment, scRNAseq was performed. We therefore sorted CD45^-^ CD31^-^ stromal cells from joints of arthritic animals receiving either no treatment or anti-TNF or anti-IL17a blocking antibodies and healthy animals (n =3 per condition). Of note, the stratification to treatment demonstrated that *Fap* expression in the clusters of lining as well as sublining fibroblasts is independent of anti-TNFα or anti-IL-17 treatment **(Fig. 2j)**. Accordingly, *Fap* expression did also not correlate with FAPI-04 uptake **(Fig. 2k)**. To exclude the biases arising from scRNAseq differential expression methods, we confirmed the ubiquitous *Fap* expression by pseudo bulk analysis **(Extended Data Fig. 2)**.

Tackling the discrepancy between *Fap* expression and FAPI-04 uptake, we studied *Fap* expression in a fibroblast subtype dependent manner. To avoid analytical bias because of unrelated exclusion of certain subtypes of fibroblasts ^13^, we integrated 33 scRNAseq datasets from various arthritis models and their controls resulting in to a total of 52.131 mesenchymal cells from joints, and performed unbiased graph-based clustering. In accordance to previous reports ^5,14–19^. The integrated dataset confirmed a high heterogeneity of joint stromal cells. Unsupervised clustering revealed seven different mesenchymal cell types **(Fig. 3a)**. There was a clear separation between lining and sublining subtypes. Furthermore, respective signature genes clearly attributed to the described sublining *Mmp3*+, *I16*+, *Pi16*+ fibroblast subtypes and robustly identified chondrocytes and osteoblasts within the dataset **(Fig. 3b)**. In the next step, similar clusters were identified in the hTNFtg197 mice challenged to *I123*mc and treated with or without neutralizing antibodies to TNF or IL-17a. Differential abundance analysis was done in order to identify significantly enriched or depleted fibroblasts subtypes. Beside overall moderate changes within the compartment of fibroblasts, a striking upregulation of so far unexplored *Cd200*+ fibroblasts was detectable upon anti-inflammatory treatment **(Fig. 3c,d)**. Those fibroblasts expressed also high levels of *Cdh11, Dkk3* but not *Lrrc15* **(Fig. 3b, Extended Data Fig. 3c)**. While the TNF blocker only slightly increased these *Cd200*+ fibroblasts, the IL-17a blocker induced a significant upregulation. In order to rule out the potential bias introduced by discrete clustering and its’ impact on cell subtype abundancies ^20^, we also implemented MELD algorithm, which assigns a condition-specific likelihood estimate to each individual cell. MELD results further confirmed that *Cd200*+ fibroblasts have a high average IL-17a inhibition associated likelihood **(Fig. 3e, Extended Data Fig. 3a)**, hence revealing their enrichment upon this treatment. Moreover, computational directed single-cell fate mapping using CellRank ^21^, when aggregating the RNA velocity estimates and transcriptional similarities, demonstrated a general increase of differentiation probability toward *Cd200*+ fibroblasts among all subsets of sublining fibroblasts upon IL-17a inhibition **(Fig. 3f, Extended Data Fig. 3b)**. This pattern of induction of *Cd200*+ fibroblasts showed strong similarity to the downregulation of FAPI-04 accumulation in the joints upon treatment. However, when we generated an additional dataset of hTNFtg197 mice having received *I12*3mc and sorted this time CD45^-^ CD31^-^ Pdpn^+^ Fap^+^ cells, integration of this dataset with the previous one showed that all FAP^+^ cells were evenly distributed across all subtypes of mesenchymal cells **(Fig. 3g, h, Extended Data Fig. 3d-h)**. Furthermore, the direct comparison between *Cd200*^+^ and *Cd200*^-^ fibroblasts did not show up with any differences of FAP expression on mRNA as well as protein level **(Fig. 3i, j)**.

**Figure 3:**
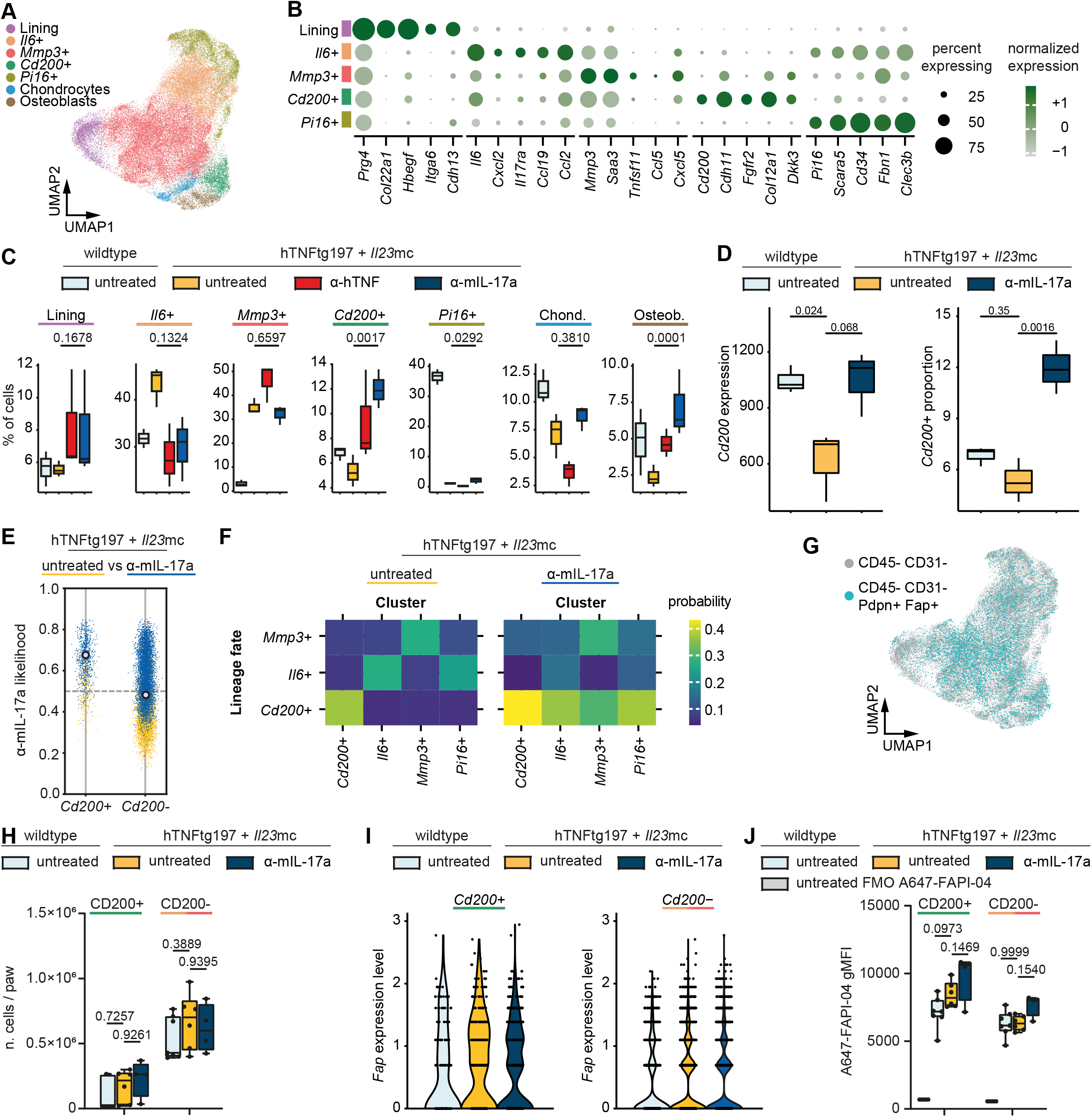
Heterogeneity of *Fap*^+^ joint stromal cells. **(a)** U-map of an integrated scRNA-seq dataset showing the heterogeneity of joint stromal cells. In total, 33 different datasets containing 52.131 CD45^-^ cells from joints of healthy controls, arthritis models and treatment/knock out variants of those models have been integrated. **(b)** Expression of the most relevant marker genes among each fibroblast cluster from the scRNA-seq dataset introduced in (a); **(c)** Differential abundancy of stromal cell populations determined by edgeR in the scRNA-seq dataset introduced in (a) containing joint stromal cells from wildtype animals and hTNFtg197 treated with Il23mc for 9 days receiving either no treatment, anti-hTNF or anti-mIL-17a treatment. Median + q + min-max are shown; N=3 per condition. **(d)** Pseudobulk-based calculation of *Cd200* expression in pooled sublining fibroblasts and differential abundance of *Cd200*^+^ fibroblasts (edgeR) in scRNA-seq dataset generated from wildtype, and Il23mc treated hTNFtg197 animals with or without anti-mIL-17a treatment. Median + quartiles + min-max are shown; N=3 per condition. **(e)** Anti-mIL-17a treatment associated relative likelihood of *Cd200*^+^ and *Cd200*^-^ sublining fibroblasts in the scRNA-seq dataset generated from Il23mc treated hTNFtg197 animals with or without anti-mIL-17a treatment determined by MELD. **(f)** Average transition probability for *Cd200*^+^, *I16*, *Mmp3*^+^ and *Pi16*^+^ sublining fibroblasts to adopt a *Mmp3*^+^, *I16*^+^ or *Cd200*^+^ terminal state using Cellrank. **(g)** U-map highlighting of *Fap*^+^ PDPN^+^ CD45^-^ CD31^-^ sorted cells from hTNFtg197 + *I123*mc animals (day 9) among the datasets shown in (a); N=5. **(h)** Flow cytometry based quantification of CD200^+^ and CD200^-^ sublining fibroblasts (PDPN^+^ PDGFRα^+^ THY1^+^CD49f) from wildtype animals and hTNFtg197 treated with Il23mc for 9 days receiving either no treatment or anti-mIL-17a treatment. Median + quartiles + min-max are shown. **(i)** *Fap* expression in *Cd200*^+^ and *Cd200*^+^ (*I16*^+^, *Mmp3*^+^) sublining fibroblasts across conditions from the scRNA-seq dataset introduced in (a). **(j)** A647-FAPI-04 geometric mean fluorescence intensity (gMFI) across CD200^+^ and CD200^-^ sublining fibroblasts (PDPN^+^ PDGFRα^+^ THY1^+^ CD49f) from wildtype animals and hTNFtg197 treated with Il23mc for 9 days receiving either no treatment or anti-mIL-17a treatment. Median + quartiles + min-max are shown.

### CD200^+^ fibroblasts sustain type 2 fate and survival of pro-resolving ILCs

As reduced FAPI-04 internalization was associated with an increase of *Cd200*^+^ fibroblasts, we next examined the functional implication of *Cd200*^+^ fibroblasts. Single sample gene set enrichment analysis of selected pathways revealed that in contrast to *Mmp3*+ and *I16*+ fibroblasts or osteoblasts, *Cd200*+ fibroblasts have strong capacity to dampen inflammatory pathways **(Fig. 4a, Extended Data Fig. 4a)**. The CD200 receptor CD200R1 showed its highest expression levels in type 2 innate lymphoid cells (ILC2s) and accordingly intercellular interaction analysis revealed strong communication probability between *Cd200*^+^ fibroblasts and ILC2s during tissue homeostasis which was lost under arthritic conditions **(Fig. 4b-d, Extended Data Fig. 4c)**. In order to further unfold the mechanism by which CD200 affects ILC2s, single cell gene co-expression network analysis was performed using hdWGCNA workflow on ILCs ^22^. Briefly this approach overcomes the unreliability of simple single cell correlations by grouping transcriptionally similar cells into meta cells, hence providing more robust correlation estimates. After identifying clusters of highly correlated gene modules, *CD200r1* appeared in the same module with prototypic ILC2 markers such as *I11r11, Gata3* and *Bc12* indicating high degree of co-expression **(Fig. 4e, Extended Data Fig. 4d, e)**. Functional enrichment analysis of genes co-expressed in this module suggested pro-survival signals upon *Cd200r1* activation **(Fig. 4f)**. Accordingly, CD200R stimulation resulted into significant reduction of cellular apoptosis of ILC2s *in vitro* **(Fig. 4g)**. Furthermore, coexpression of *Cd200r1* and *Gata3* suggested a potential role of CD200R signaling in maintaining the ILC2 phenotype, therefore we studied this potential role by computationally knocking out the *Cd200r1* on ILCs. To this end we utilized the dynamo framework that provides an analytical solution to velocity vector fields ^23^, hence enabling the study of genetic perturbation on cell differentiation. Indeed, the in silico knock out of *Cd200r1* predicted a cell fate redirection towards an ILC3-like fate **(Fig. 4h)**. Appropriately, the treatment of arthritic mice with CD200-Fc was associated with significant reduced inflammation and FAPI-04 uptake in the joints and resulted into significant reduced joint damage **(Fig. 4i)**.

**Figure 4:**
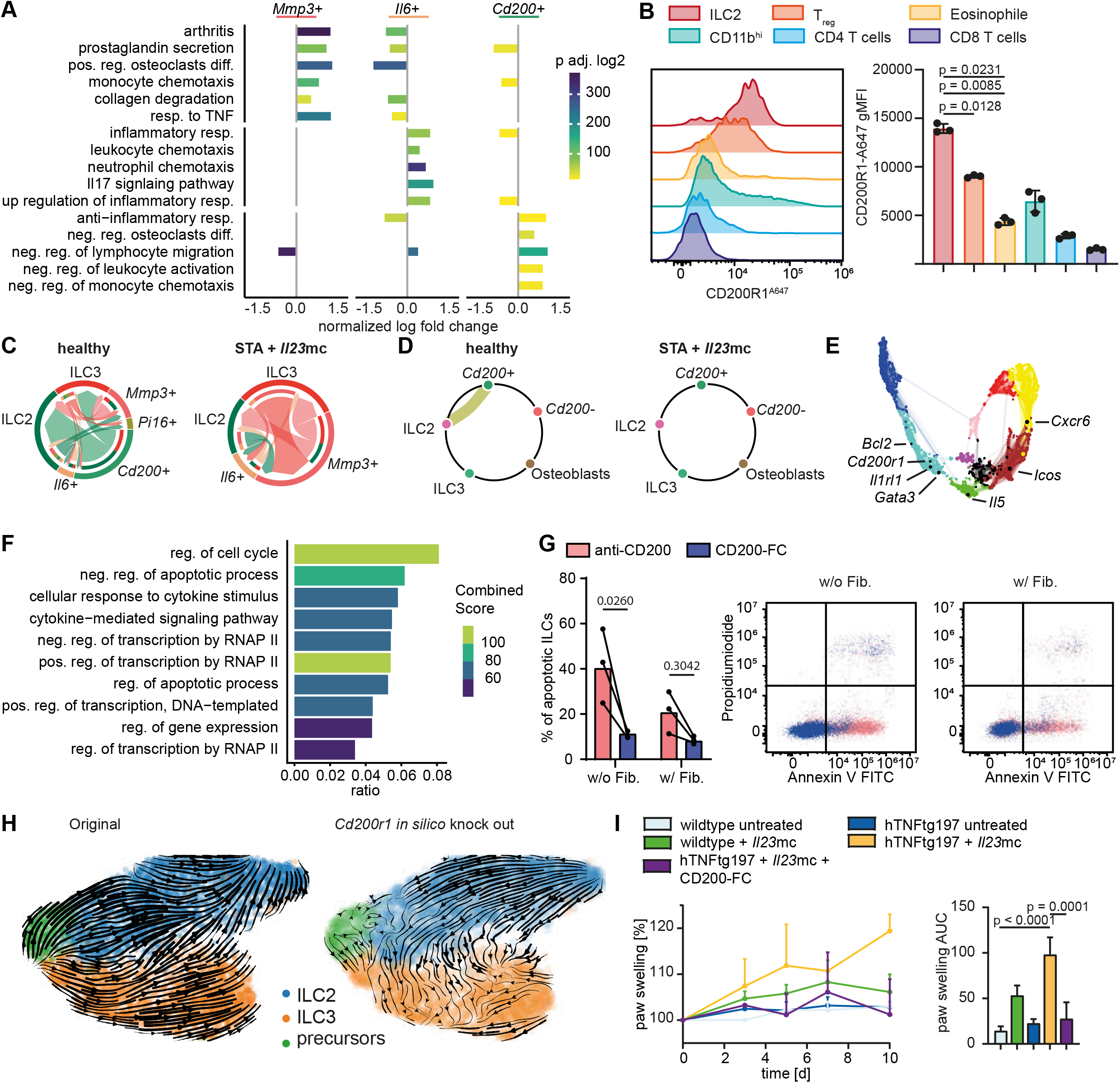
*Cd200*^+^ fibroblasts sustain type 2 fate and survival of pro-resolving ILCs. (**a)** Log-fold changes of the Enrichment scores for relevant inflammatory, anti-inflammatory, and damage related GO and KEGG terms using ssGSEA. Wilcoxon Rank Sum test was used to find significantly negatively or positively enriched terms in each cluster (adj. p-value < 0.05; LFC > 0.25). **(b)** Fluorescence intensity of CD200R1 (Ox-110) among pro- and antiinflammatory cells found in the inflamed synovial paw joint of hTNFtg197 + Il23mc animals. Mean + SD is shown. **(c)** Chord diagram of the communication probability of each cluster in the combined dataset of fibroblasts and ILC2s from either healthy (HC) or KBxN serum transfer arthritis model (STA) treated with *I123*mc for 9 days. Shown are the >80% enriched ligand receptor pairs for each condition. Fibroblasts from this model show a similar pattern compared to hTNFtg + *I123*mc derived fibroblasts (Extended Data Fig. 4b) **(d)** Chord diagram of only the CD200-CD200R communication probability from (c). **(e)** Modules of highly correlated genes from the hdWGCNA analysis of R5+ ILCs sorted from the ankle joints of either healthy or STA + *I123*mc animals. *Cd200r1* appears in the same module (cyan) with prototypic ILC2 markers such as *I11r11, Gata3* and *Bc12*. **(f)** In the cyan module of (e) enriched “GO biological processes”-terms using enrichR. **(g)** Apoptosis rate of murine ILC2s determined by Annexin-V/propidium iodide staining after culture in presence or absence of chimeric CD200-Fc and murine synovial fibroblasts. **(h)** Prediction of the cell fate redirection after the in silico knock out of *Cd200r1* using dynamo. **(i)** Paw swelling of wildtype, wildtype + *I123*mc, hTNFtg197, hTNFtg197 + *I123*mc and hTNFtg197 + *I123*mc + CD200-Fc animals. Mean + SD are shown; N = 3, 4, 3, 3, 4, respectively.

### FAPI uptake regulated by receptor internalization

Beside hallmark features of resolution of inflammation attributed to *Cd200*^+^ fibroblasts, gene set enrichment analysis revealed a significant difference in the receptor internalization capacities upon treatment with IL-17a blocking antibodies **(Fig. 5a, b)**. To validate these observations, synovial fibroblasts were cultured under endocytosis affording and blocking conditions in presence of fluorescently labeled A647-FAPI-04. Blocking endocytosis by addition of 450 mM sucrose or decreasing the temperature to 4°C significantly decreased the geometric mean fluorescence intensity **(Fig. 5c)**. Also in arthritic mice, A647-FAPI-04 uptake was significantly reduced upon treatment with IL-17 inhibitors **(Fig. 5d)**. Thus, FAPI-04 signal intensity as assessed by PET does not correlate with FAP expression levels on the surface of total synovial fibroblasts but rather reflects the functionally active receptor internalization capacities of fibroblasts via FAP within the mesenchymal compartment. FAP related receptor internalization of FAPI-04 dropped with the increase of *Cd200*^+^ fibroblasts and their induction of resolution of inflammation upon treatment with IL-17a blocking antibody. To further stratify the relationship between FAP related receptor internalization capacity and joint damage, we evaluated the correlations between both parameters. Indeed, FAP related receptor internalization strikingly correlated with joint damage **(Fig. 5e)**. *Vice versa,* FAP related receptor internalization significantly decreased with the upcoming *Cd200*^+^ related pro-resolving capacity in a correlative manner.

**Figure 5:**
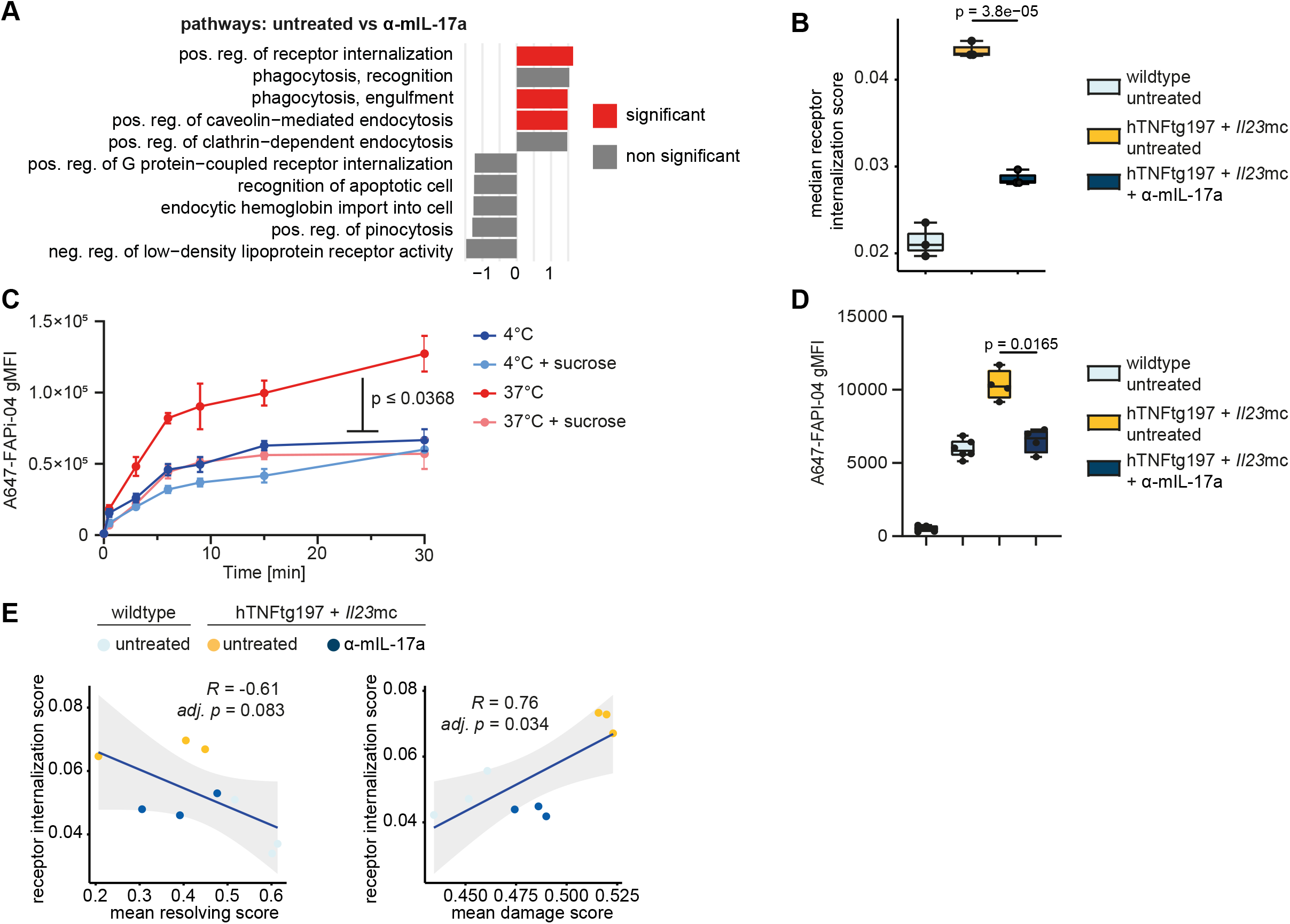
FAP-related receptor internalization differs between clusters of synovial fibroblasts. **(a)** Gene set enrichment analysis using fgsea for all terms of the “endocytosis” (GO:0006897) set based on pseudobulk determined DEGs of *Fap*-expressing clusters between untreated and anti-mIL-17a treated hTNFtg197 + *I123*mc conditions. Top five positively and negatively enriched terms are shown. **(b)** Receptor internalisation score across *Fap*-expressing clusters between conditions. The leading edge gene list of the “pos. reg. of receptor internalization” (GO:0002092) was used to calculate the score for each cell using UCell and was summarized over all cells for each replicate. Median + quartiles + min-max are shown. **(c)** A647-FAPI-04 gMFI over time in cultured murine joint fibroblasts under optimal (37°C) and endocytosis blocking conditions (37°C + sucrose, 4°C, 4°C + sucrose). Mean + SD are shown; N= 5. **(d)** A647-FAPI-04 gMFI across CD200^+^ and CD200^-^ sublining fibroblasts (PDPN^+^ PDGFRα^+^ THY1^+^ CD49f) from wildtype animals and hTNFtg197 treated with Il23mc for 9 days receiving either no treatment or anti-mIL-17a treatment. A647-FAPI-04 was injected i.v. 24h prior to sacrifice. Mean + SD are shown. **(e)** Pearson correlation between FAPI-04 internalization and joint damage or pro-resolving parameters respectively.

### Conserved phenotype of pro-resolving CD200^+^ DKK3^+^ in mice and humans

Finally, we examined the potential translation of the functional relationship between joint damage and FAPI-04 uptake, and the reduction of FAPI-04 accumulation during cellular shift in the mesenchymal compartment to pro-resolving CD200^+^ fibroblasts in humans. Therefore, we first created a comprehensive map of human PsA synovial single cells by integrating four publicly available datasets ^16,24^ and identified the major cellular heterogeneity **(Fig. 6a)**. This approach was followed by another integration step in which murine *I123*mc treated hTNFtg197 fibroblasts were mapped to their human counterpart, which led to the identification of similar subtypes of fibroblasts in human, forming distinct transcriptional neighborhoods, observable on the U-map, as well as with similar markers expression **(Fig. 6b-d, Extended Data Fig. 5a)**. To study how each of the fibroblast’s subtypes are spatially distributed in the tissue, and to quantify their colocalization with different immune cells, we then performed spatial transcriptomic analyses of synovial biopsies from patients with psoriatic arthritis in the stage of active disease **(Fig. 6e, Extended Data Fig. 5b)**. The inherent low spot resolution, and the sparsity of spatial transcriptomic data encouraged us to employ a resolution enhancement method implemented by BayesSpace algorithm ^25^. This strategy assisted us to identify spatially heterogeneous clusters on the tissue with fine resolution, corresponding to the pathogenic and anatomical regions from the Visium slide **(Fig. 6e)**. Scoring of the gene modules corresponding to various cell types, followed by a hierarchical clustering of these scores across the spatial clusters, revealed the co-occurrence of CD200^+^/DKK3^+^ fibroblasts with pro-resolving immune cells such as ILC2s and eosinophils in regions with no apparent inflammation **(Fig. 6f)**. In contrast, MMP3^+^/IL6^+^ fibroblasts co-localized with inflammatory immune cells in regions with active inflammatory phenotype. Moreover, scoring the pathways previously introduced in the murine fibroblast subtypes confirmed the inflammatory signature in regions where MMP3^+^/IL6^+^ clusters reside, and an anti-inflammatory signature in regions where CD200^+^/DKK3^+^ are located **(Fig. 6g)**.

**Figure 6:**
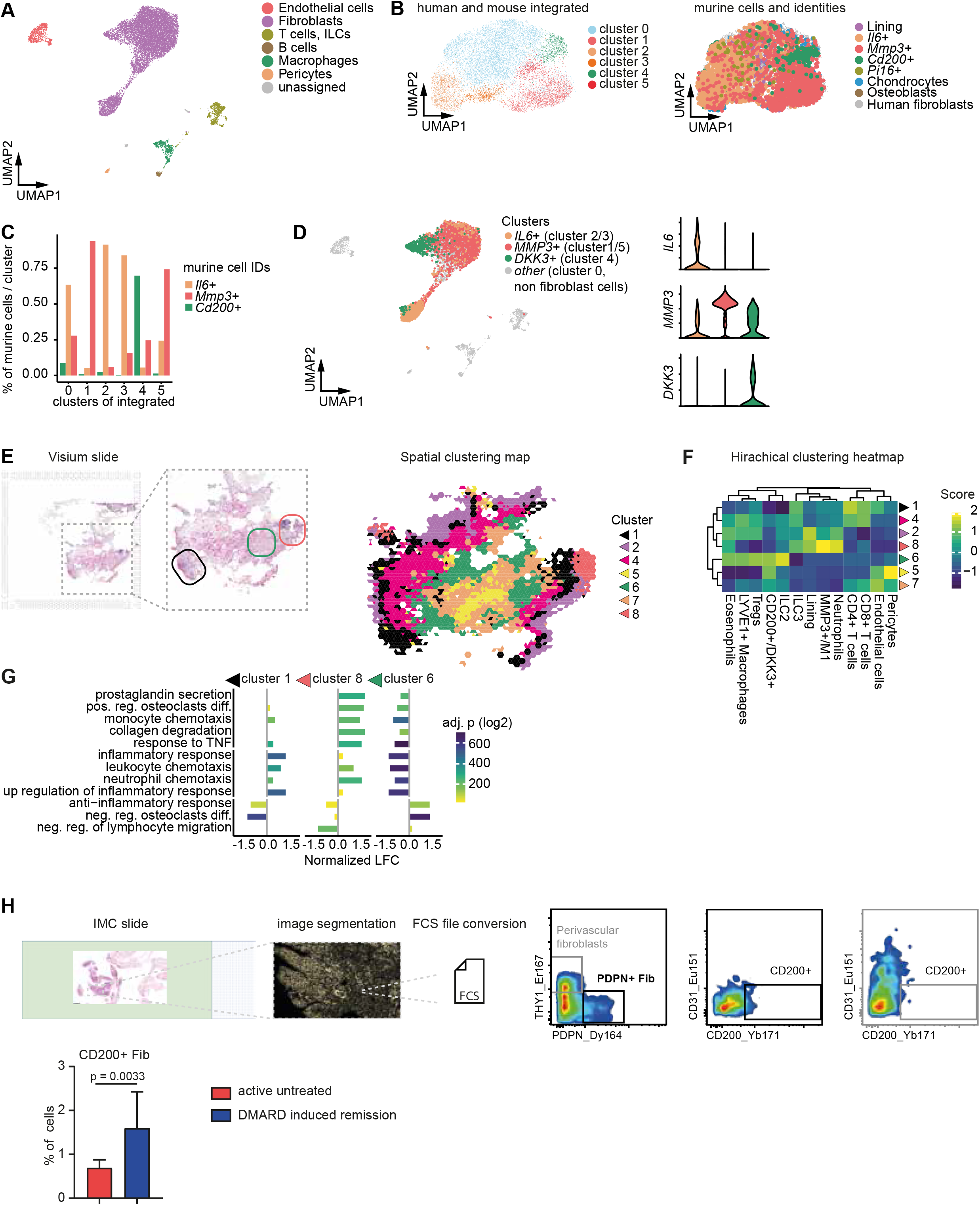
Translation to human synovium of patients with psoriatic arthritis. **(a)** U-map of an integrated scRNAseq dataset showing the heterogeneity of human synovial cells of PsA patients. In total, four different datasets containing 11,414 cells from two CD45^+^ sorted single cell datasets, and two whole tissue single cell datasets were integrated. **(b)** U-map of fibroblasts of integrated human PsA patients and Il23mc treated hTNFtg197 mice showing the unsupervised identified clusters (left) and the previously identified murine fibroblast subsets (right). **(c)** Bar plot quantifying the percentage of *Cd200*^+^, *I16*^+^ and *Mmp3*^+^ murine cells in each of the unsupervised identified clusters. **(d)** U-map highlighting the previously in the integrated dataset identified clusters (a), forming transcriptionally distinct clusters on the U-map (left). Violin plot showing the expression of the top markers of these clusters (right). **(e)** Representative visium slide of human synovial tissue from a PsA patient and visualization of the identified spatial clusters. Regions of interests are highlighted (left), colors correspond to the identified spatial clusters. BayesSpace algorithm was implemented to enhance the spot resolutions and identify the spatially heterogeneous regions. **(f)** Hierarchically clustered heatmap of cell type specific gene module scores across the spatial clusters, calculated with UCell on the BayesSpace assisted enhanced gene expression. **(g)** Log-fold changes of pathways scores previously introduced in murine fibroblasts subtypes for spatial clusters 1 and 8 (IL6^+^ and MMP3^+^ cells), and 6 (CD200^+^ DKK3^+^). **(h)** Imaging mass cytometry (IMC) analysis workflow for the detection of CD200^+^ fibroblasts in human synovial biopsies. 6 PsA patients, 3 with untreated active disease, 3 with a DMARD induced remission were analyzed.

Finally we employed imaging mass cytometry (IMC) to disentangle CD200^+^ fibroblasts in the synovium of PsA patients. In line with the spatial transcriptomic analysis and the murine data, we observed a significantly increased proportion of CD200^+^ fibroblasts in the synovium of PsA patients in remission after successful DMARD treatment **(Fig. 6h)**.

## DISCUSSION

Our data demonstrate a shift in the fibroblast compartment during successful initiation of resolution of inflammation and protection from joint damage. This phenotypic switch protects from joint damage and can be visualized by PET/CT. The correlation between FAPI-04 uptake and presence of functionally distinct fibroblast subtypes can be used not only to visualize active joint damage but might also improve drug monitoring and silent joint destruction under DMARD therapies in particular in patients with disease flares and persistent inflammatory disease activity^26^.

FAPI-04 are quinolone based FAP inhibitors that can be linked to radioactive tracers but also other dyes. Therefore, PET/CT might be not the only imaging modality that enables the use of FAPIs as diagnostic tool to visualize the fibroblast compartment and might provide the possibility to image tissue damage in a diagnostic surrounding without radiation^10^.

Effects of currently used DMARDs upon phenotypic changes of fibroblast subtypes and their contribution to resolution of inflammation are not investigated so far. Herein, we show effects of both, TNF and IL-17A blockers to induce CD200^+^ inflammation-resolving fibroblasts with their capacity to protect from joint damage. While TNF inhibitors exhibited mild effects in this study, IL-17A blockers significantly induced CD200^+^ fibroblasts and mostly abolished FAPI uptake in mice and humans. These observations have to be further evaluated in clinical trials with different patient cohorts.

Furthermore, the data demonstrate that the microenvironment has a major impact on functionality of immune cells that induce resolution of inflammation. Studies so far describe fibroblast characteristics to induce inflammation and tissue damage^5,14,27^. Here we show, that resolution of inflammation is not only associated with a reduced abundancy of such inflammatory and destructive fibroblasts but rather with an active change of the composition of the fibroblast compartment. Indeed destructive subtypes loose cellular abundancy and functionality. Beside these effects negatively influencing fibroblast activity, the data show that initially destructive fibroblasts transdifferentiate into subtypes with opposing functions with features to resolve inflammation. These functional properties mainly target ILC2s that are well described mediators of resolution of inflammation in arthritis^11^. CD200R signaling leads to stabilization of the ILC2 fate and decrease transdifferentiation into exILC2/ILC3 cells. This shift in the ILC balance is an important mechanism to inhibit inflammation by release of inflammatory cytokines on the one hand, and in addition fostering Treg activation and increase of eosinophilic granulocytes^3^. Therapies targeting fibroblasts are not available for treatment of chronic inflammatory diseases^6,27^. CD200 however showed strong effects to control inflammation and tissue damage. Targeting of CD200 might thus provide a novel therapeutic option to efficiently but also safely interfere with tissue damage and allow to restore tissue homeostasis in chronic inflammatory joint diseases.

## Acknowledgements

The authors thank to Dr. Melanie Rose and Jianli Tu for excellent technical assistance.

## Author Contributions

Design of the study: SR, HM, CS, GS, AR; Acquisition of data: SR, HM, CS, AA, AS, SM, CT, MA, MGR, CX, KTY, LL, HL, EK, JK, AR; Interpretation of data: SR, HM, CS, AA, AS, SM, CT, HM, TK, OP, JC, GS, AR; Support of material: SM, JHWD, TB, TK, OP, JC, GS; Manuscript preparation: SR, HM, GS, AR

---

**Extended Data Fig. 1:**
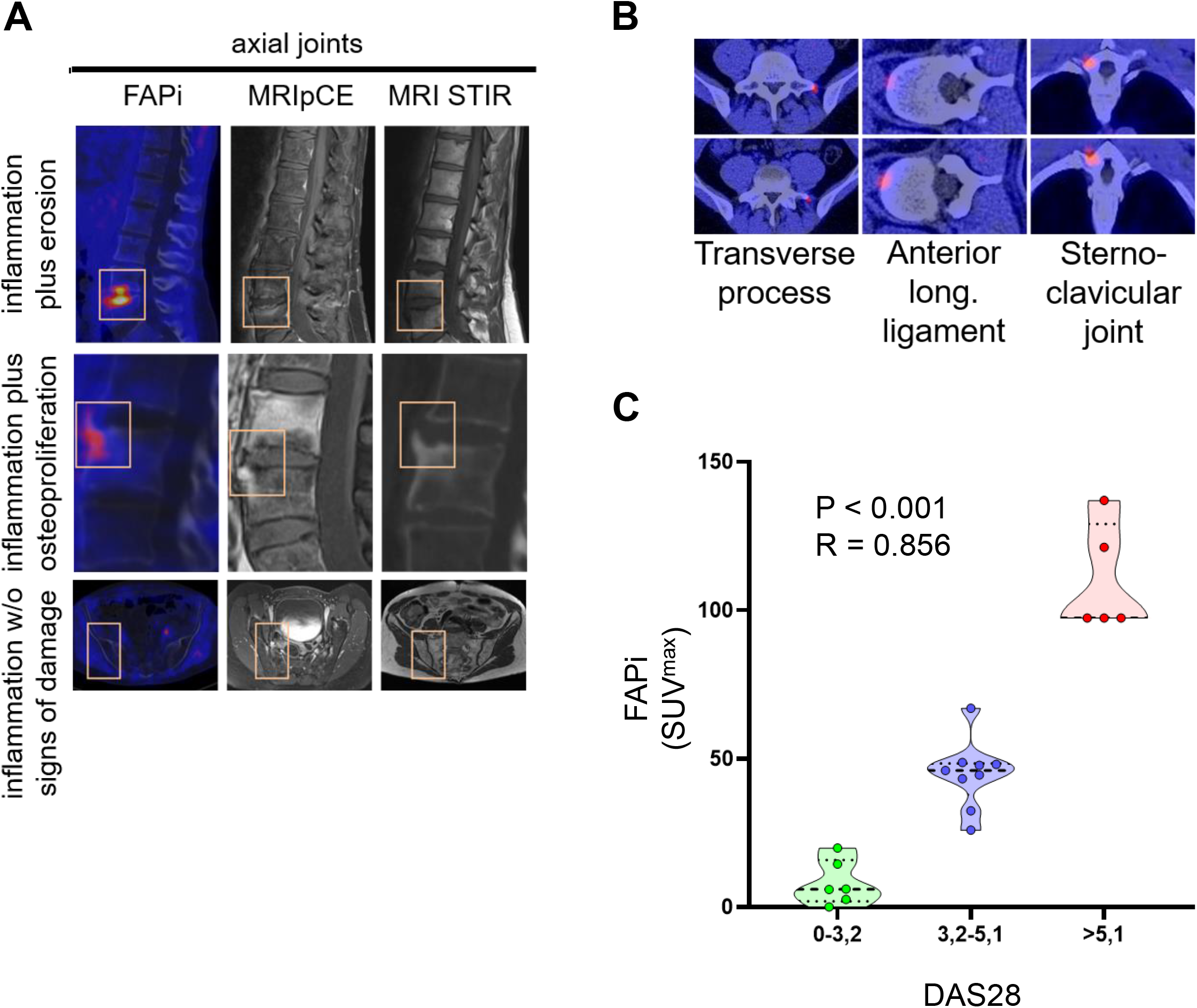
**(a)** Comparative analysis of axial ^68^Ga-FAPI-04 PET/CT and MRI scans from the same patients; **(b)** representative examples of different localizations of ^68^Ga-FAPI-04 uptake at axial joints as assessed by PET/CT **(c)** Pearson correlation analysis of ^68^Ga-FAPI-04 uptake (maximum standardized uptake value; SUV^max^) and composite score DAS28 (disease acitivity score of 28 joints in patients with rheumatoid arthritis).

**Extended Data Fig. 2:**
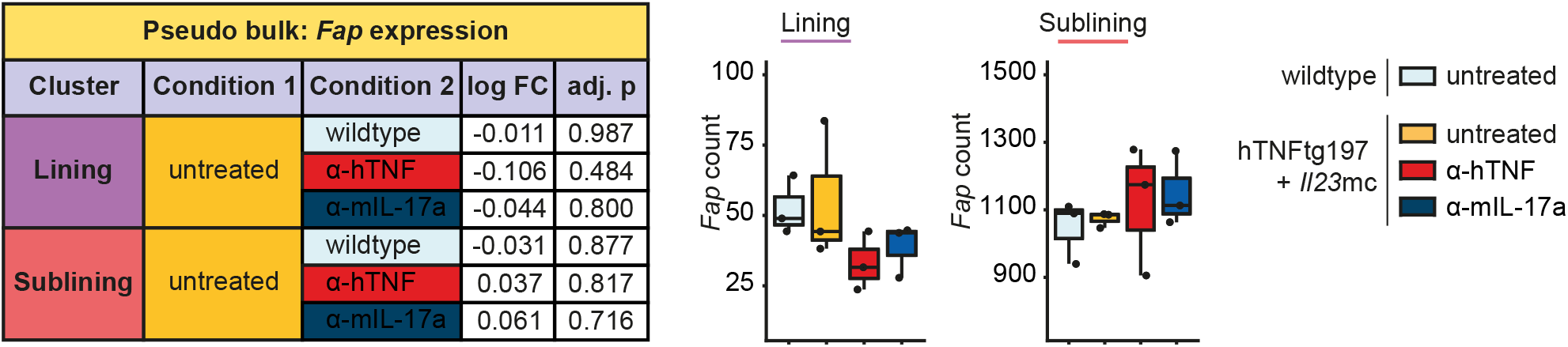
Pseudobulk-based calculation of *Fap* expression in lining and sublining fibroblasts of a scRNA-seq dataset generated from sorted fibroblasts of wildtype animals and hTNFtg197 treated with *I123*mc for 9 days receiving either no treatment, anti-hTNF or anti-mIL-17a treatment.

**Extended Data Fig. 3:**
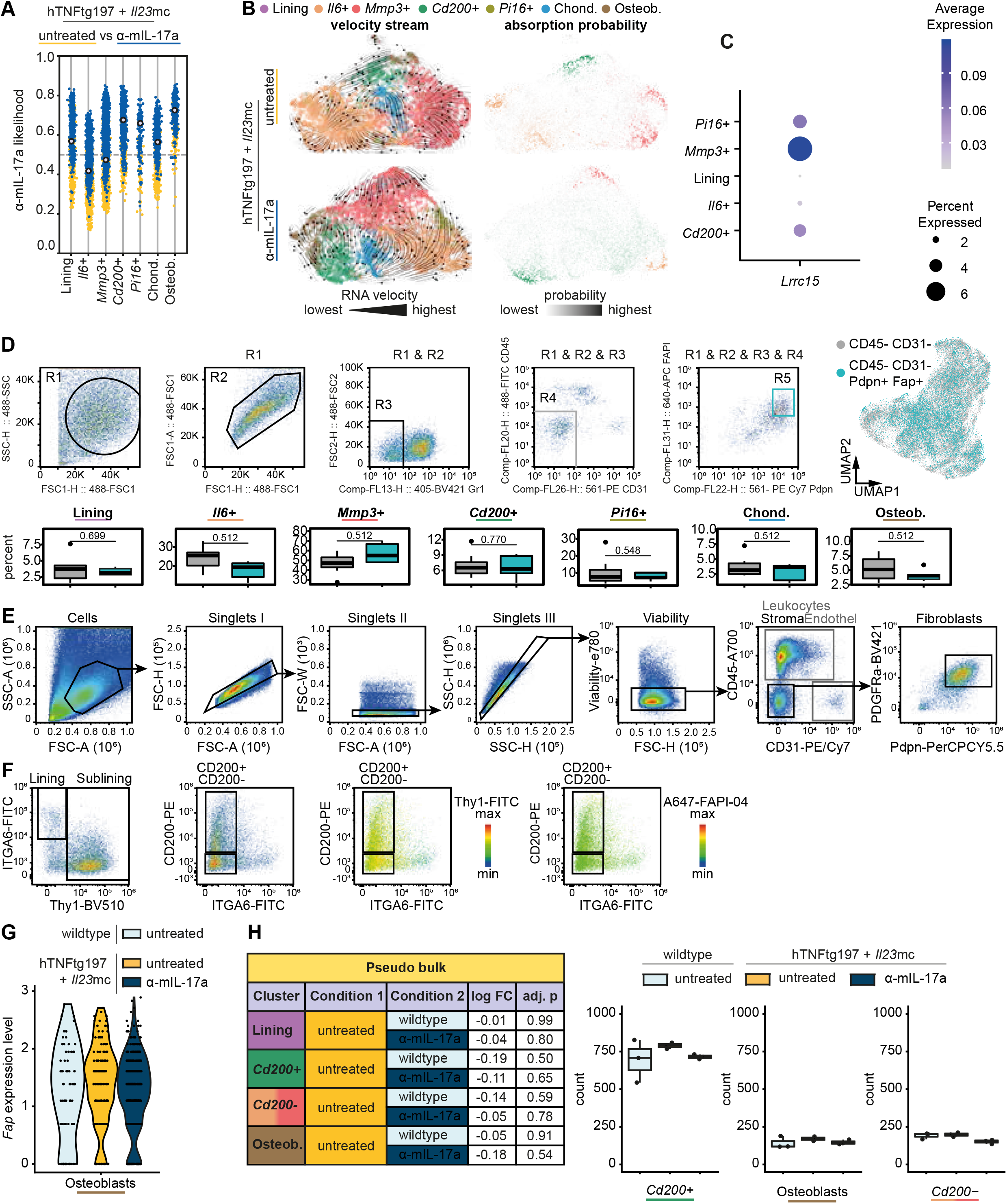
**(a)** Anti-mIL-17a treatment associated relative likelihood of each fibroblast cluster in the scRNA-seq dataset generated from Il23mc treated hTNFtg197 animals with or without anti-mIL-17a treatment determined by MELD. **(b)** Velocity streams and absorption probabilities of terminal states within the scRNA-seq dataset generated from *I123*mc treated hTNFtg197 animals with or without anti-mIL-17a are visualized over the U-map. **(c)** Expression of *Lrrc15* across sublining fibroblast clusters. **(d)** Gating strategy to sort Fap^+^ Pdpn+ stromal cells from joints for scRNA-seq. FCS file for representation is directly derived from the sorter. Abundancy of fibroblast clusters between CD45-CD31- and CD45-CD31+ Fap+ Pdpn^+^ stromal cells. Median + quartiles are shown. **(e)** Gating strategy to identify synovial fibroblasts in joints of arthritic mice. **(f)** Markers for lining (ITGA6^+^) and sublining (Thy1^+^) fibroblasts expressed on synovial fibroblasts in mice. Strategy to identify CD200^+^ and CD200^-^ synovial sublining fibroblasts. Heatmaps show expression of Thy1 and Fap among synovial fibroblasts. **(g)** *Fap* expression across conditions for osteoblasts. **(h)** Pseudobulk-based calculation of *Fap* expression on lining and sublining fibroblasts of a scRNA-seq dataset generated from sorted fibroblasts of wildtype animals and hTNFtg197 treated with *I123*mc for 9 days receiving either no treatment, anti-hTNF treatment or anti-mIL-17a treatment.

**Extended Data Fig. 4:**
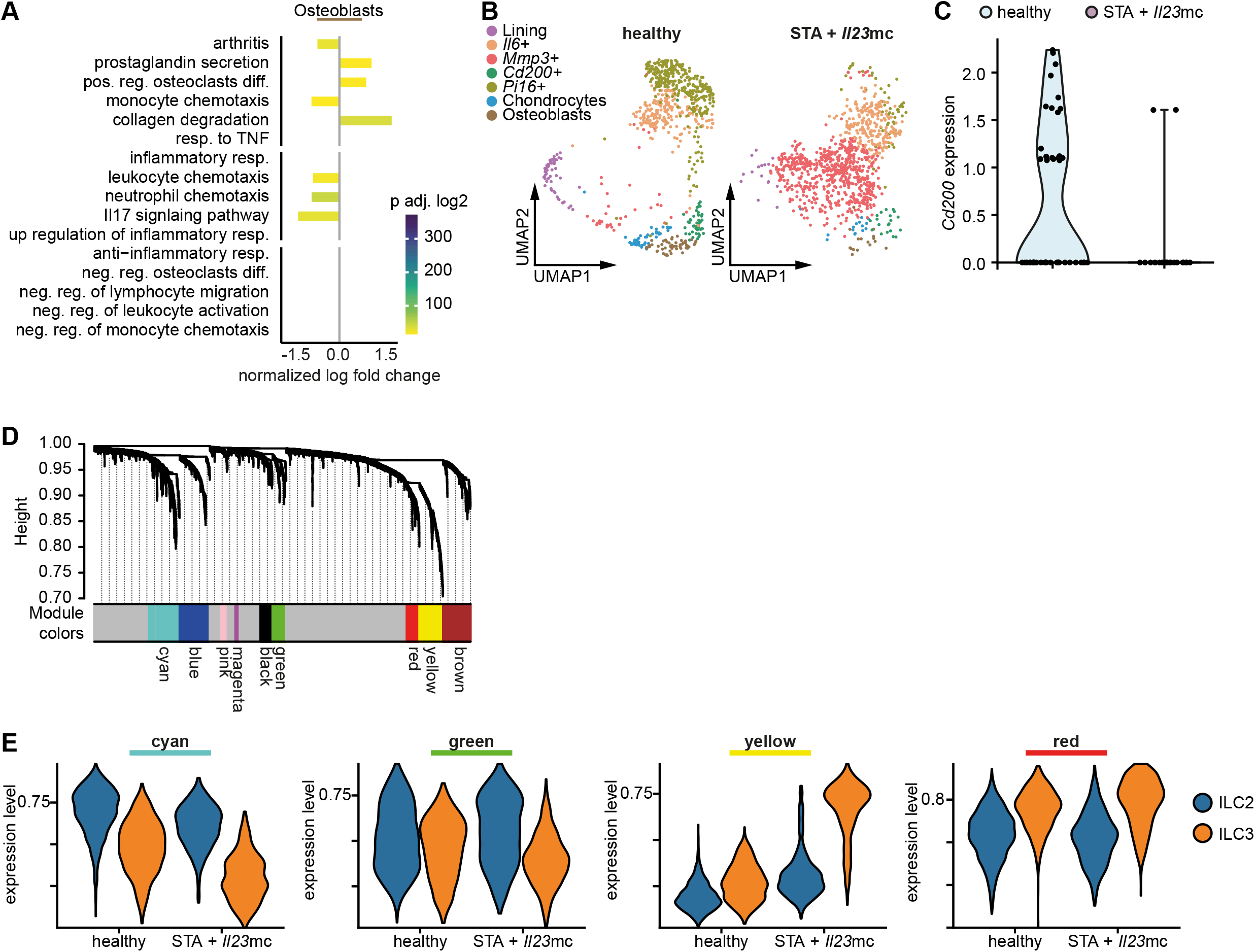
**(a)** Log-fold changes of the enrichment scores for relevant inflammatory, anti-inflammatory, and damage related GO and KEGG terms using ssGSEA shown for osteoblasts. Wilcoxon Rank Sum test is used to find significantly negatively or positively enriched terms in each cluster (adj. p-value < 0.05; LFC > 0.25). **(b)** U-map of fibroblasts from healthy controls and STA + *I123*mc model, both sorted from R5-fate mapping (ILC2 reporter) animals. **(c)** Violin plots of *Cd200* expression among healthy controls and STA + *I123*mc model from the R5-fate mapping animals. **(d)** Representation of highly correlated gene modules. Gene co-expression networks were calculated with hdWGCNA, tree-cut algorithm was used to identify highly correlated gene modules (colored), grey sections are non-correlated genes. **(e)** Scores for each module of cp-expressing genes stratified by condition and cell type calculated by UCell.

**Extended Data Fig. 5:**
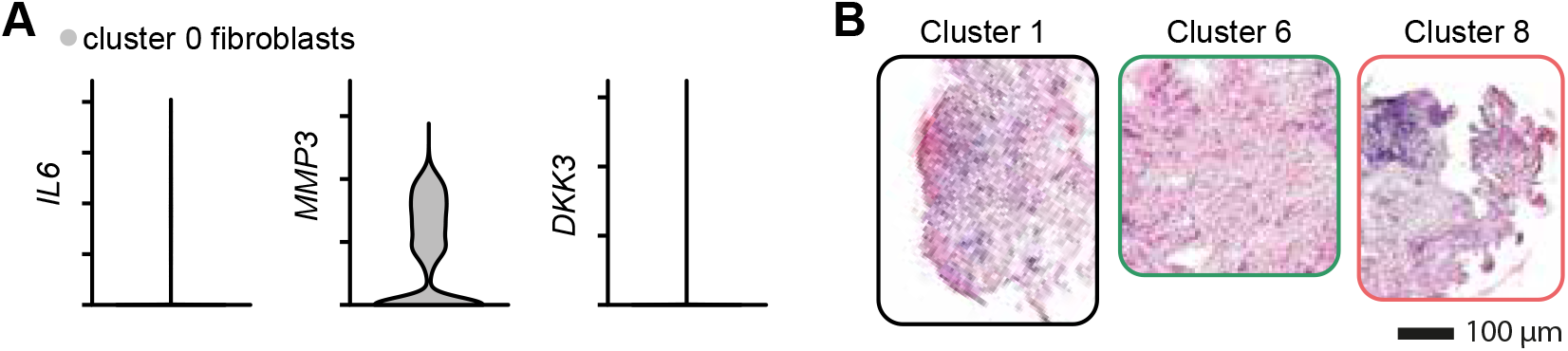
**(a)** Violin plot showing the expression of the top markers as in Fig. 6D for cluster 0 containing mixed mouse IDs and mostly human cells. **(b)** Higher magnifications of the ROIs identified in Fig. 4e.

## Material and Methods

### Experimental approaches

Experiments were not performed in a blinded fashion except when specifically indicated. There were no exclusion criteria for the human and animal experiments. Mice were stratified according to sex and then randomized into the different groups. Cells from human donors were also randomized.

### Patient characteristics

One hundred forty eight patients underwent ^68^Ga-FAP inhibitors (FAPI)-04 PET/CT. All patients underwent the examinations as part of the clinical workup in order to accumulate diagnostic evidence and potentially optimise their individual treatment. ^68^Ga-FAPI-04 is an investigational radiopharmaceutical in arthritis and not yet approved by the Food and Drug Administration or the European Medicines Agency. It was, therefore, administered under the conditions outlined in §13 (2b) of the Arzneimittelgesetz (German Medicinal Products Act) and in compliance with the Declaration of Helsinki. Written informed consent for evaluation and publication of their anonymised data was obtained from all patients. Subjects with RA (n = 20) fulfilled the 2010 American College of Rheumatology (ACR) classification criteria for RA ^1^. Subjects with PsA (n = 85) fulfilled the Classification Criteria for Psoriatic Arthritis (CASPAR) ^2^. Subjects with axial spondylitis (n = 28) fulfilled the ASAS criteria ^3^. Fifteen non-arthritic patients served as controls. There was no significant difference in sex and age distribution between the groups. Disease activity was measured by disease related composite scores as outlined in the figure legend. For synovial tissue analyses samples from untreated patients with active PsA (n = 4) and patients with PsA in remission receiving treatment with disease-modifying antirheumatic drugs (TNF inhibitors, n = 4) were analyzed. Samples were retrieved by synovial biopsy in centers from Barcelona, Spain, and Erlangen, Germany.

### Animals

hTNFtg197 (B6.Cg-Tg(TNF)197Gkl) animals have been described in ^4^. hTNFtg197 have been maintained on a C57Bl/6N background. R5 mice (B6(C)-*I15<tm1.1(icre)Lky>*) have been described in ^5^. They have been crossed onto R26^tdTomato^ (B6.Cg-*Gt(ROSA)26Sor<tm9(CAG-tdTomato)Hze>*) reporter lines. From the resulting R5 x R26^tdTomato^ mice, only animals with heterozygous wt/mut *I15* locus were used because *I15* mut/mut animals display a severe arthritis phenotype ^6^. All animals were kept under SPF conditions.

### Models of inflammatory arthritis

STA was induced in R5 x R26^tdTomato^ mice by intraperitoneal injection of 200μl arthritogenic serum from K/BxN mice ^7^. Swelling of front and hind paws as well as ankles was monitored using callipers, averaged and reported as the change from baseline. Grip strength was determined by evaluating the resistance of the mice to release from the cage grid: 3 (full grip strength), 2 (slightly reduced grip strength), 1 (severely reduced grip strength), 0 (no grasping reflex).

hTNFtg197 animals were scored from 8-9 weeks of age for 9 days for symptoms of arthritis: Swelling of hind paws and ankles was monitored using callipers, averaged and reported as the change from baseline. Grip strength was determined by evaluating the resistance of the mice to release from the cage grid: 3 (full grip strength), 2 (slightly reduced grip strength), 1 (severely reduced grip strength), 0 (no grasping reflex).

*I123*mc treatment was performed together with serum transfer or on the first day of scoring of hTNFtg197 or their respective healthy littermates. The *I123*mc vector, which encodes, under control of the albumin promoter for efficient expression in hepatocytes, led by a secretion sequence for the efficient release into the blood stream, for the two subunits IL-12b (IL-12p40) and IL-23a (IL-23p19) linked by a flexible region, was a kind gift from Stefan Wirtz (Department of Internal Medicine 1, Erlangen, Germany). The vector was produced in *Escherichia coli* DH5α grown in terrific broth medium without removing the vector backbone. Plasmids were purified using the PureLink HiPure Plasmid Maxiprep Kit (Thermo Fisher) followed by the MiraCLEAN Endotoxin Removal Kit (Mirus) to ensure efficient removal of endotoxin. 3 μg of naked plasmid DNA in Ringer’s solution with a volume equivalent to 10 % of body weight was administered by hydrodynamic gene transfer into the tail vein ^8, 9^.

Antibody treatment was started on day 1 after hydrodynamic gene transfer. Animals received intraperitoneally 100μg of anti-hTNF (Infliximab, MSD) or anti-mIL-17a (17F3) on day 1 and day 7.

### PET imaging

Mice were laid on a heating pad (37 °C) and were anesthetized with O_2_/isoflurane (3 % isoflurane, 1.2 L/min O_2_). A venous access was laid into the tail vein and mice were transferred to the Inveon PET scanner (Siemens). A dynamic PET scan was started from 0 to 60 min after injection of 2 MBq of ^68^Ga-FAPI-04 in 100 mL of isotonic saline solution. For blocking, 80 nmol FAPi-04 derivate alkyne 11 ^10^ were co-injected. After iterative maximum a posteriori image reconstruction of the decay- and attenuation-corrected images, regions of interest were defined using PMOD (PMOD Technologies). The mean radioactivity concentration within the regions of interest was converted to %ID/g and the AUC of signal over time was calculated and normalized to the mean AUC of a corresponding blocking group resulting in the specific uptake to correct for group-based cofounders.

### Histological processing of murine samples

Joints were fixed in PBS containing 4 % (w/v) formaldehyde for 16 - 24 h. Bone tissue was decalcified in 0.5 M EDTA pH 7.4 before embedding and cutting. Tissue was dehydrated and infiltrated with paraffin, embedded in paraffin blocks, cut to 1 μm thin sections and mounted on slides. Paraffin embedded thin sections were deparaffinized by heating the slides for 30 min at 65 °C and washing in Histo Clear (National Diagnostics) and rehydrated with a series of 100 % (v/v) ethanol, 95% (v/v) ethanol, 80 % (v/v) ethanol, 60 % (v/v) ethanol, water. Mayer’s Haematoxylin solution (Merck) was applied for 10 min. Haematoxylin was blued by rinsing the slides for 10 min in tap water. Eosin counter stain (0.3 % (w/v) Eosin Y (Sigma), 0.01 % (v/v) acetic acid) was applied for 3 min and washed off with deionized water. Tissue was dehydrated in isopropanol, covered with Roti Mount mounting medium (Carl Roth) and a cover slip. Slides were digitalized on a NanoZoomer S60 (Hamamatsu) and ROIs exported as TIFF files. The quantification of the cell nuclei was done using the Fiji ^11^ function “Colour Deconvolution” with “H&E” vector, followed by “Auto Threshold” function with “Otsu dark” vector for deconvoluted channel 1 to convert into a binary image. After removing noise and “Watershed” separation, the segmented nuclei were counted using the “Analyze particles” function.

### Micro-CT

Calcaneus and tarsal bones were scanned by X-ray micro computed tomography (micro-CT) on a μCT 35 (Scanco Medical). The acquisition parameters were as follows: voltage 40 kV, x-ray current 250 μA, exposure time 5000 ms/projection, matrix 1024 × 1024 pixel and voxel size in reconstructed image 9 μm. The number of projections was dependant on the bone analysed. Erosion/osteoproliferation was assessed by 3 independent blinded assessors and scored: 0 (no erosion/no osteoproliferation); 1 (negative roughness/positive roughness); 2 (pitting/bulging); 3 (full thickness holes/bone spurs).

### Mouse tissue digestion

Bones with intact ankle/wrist joints were dissected. Tendons and joint capsules were cut. Tissue was digested in RPMI 1640 containing 1.25 mg/ml collagenase D (Roche), 0.2 mg/ml of Dispase II (Sigma-Aldrich) and 0.1 mg/ml of DNase I (Roche). Samples were incubated for three times at 37 °C for 20 min and constant shaking (2,000 rpm). Every 20 min supernatant was collected, filtered (< 70 μm) and fresh digestion medium was added. RBC lysis was performed using RBC lysis buffer (Biolegend). Resulting cell suspensions were filtered (< 40 μm) and cell counts were obtained after trypan blue staining using an automated cell counter (BioRad).

### Flow cytometry and cell sorting

Cells were stained on ice for 30 min in PBS containing 2 % (v/v) FBS and 5 mM EDTA. Dead cells were excluded using Zombie Violet (Biolegend) or Fixable Viability Dye eFluor 780 (Thermo Fisher). Antibodies used to analyse/sort fibroblasts were anti-CD45 (30-F11), anti-CD31 (390), anti-Podoplanin (8.1.1), anti-PDGFRa (APA5), anti-THY1 (30-H12), anti-CD49f (GoH3), anti-CD200 (OX-90), Alexa Fluor 647 coupled FAPi-04. Antibodies used to analyse CD200R1 expression were: anti-CD200R1 (OX-110), anti-CD45 (30-F11), anti-Siglec F (S17007L), anti-CD3e (145-2C11), anti-CD4 (RM4-5), anti-CD11b (M1/70), anti-CD25 (PC61), anti-CD90.2 (30-H12). Antibodies used to sort R5+ ILCs and Fibroblasts: GR-1 (RB6-8C5), anti-CD45 (30-F11), CD3e (245-2C11), anti-TCR beta (H57-597), anti-TCR gamma/delta (GL3), anti-CD11b (M1/70). All antibodies were from Biolegend. Samples were acquired using a Beckman Coulter Gallios (*ex vivo* samples) or a Beckman Coulter Cyotflex S (*in vitro* samples) and analysed by FlowJo, v.10.8. Cell sorting was performed using a Beckman Coulter MoFlo Astrios EQ. Sorted populations were reanalysed to determine target cell purity post sorting (> 98% purity).

### Generation and sequencing of droplet-based single cell RNA sequencing libraries

Different single cell RNA sequencing libraries of synovial cells have been generated (**Suppl. Tab. 1**). Cells were sorted and purified cells were stained on ice for 30 min in PBS containing 1 % (w/v) BSA with hashtag antibodies (TotalSeq-B0301/B0302/B0303/B0304B0305 anti-mouse Hashtag 1/2/3/4/5, respectively, Biolegend). Cells were washed, counted and concentrated to 1,000 cells / μl prior to pooling. Up to 25,000 cells (hyper-loading) pooled from 3-5 animals were loaded into a single well of a Chromium chip G (10x Genomics). 3’ gene expression libraries were generated using Chromium Next GEM Single Cell 3’ Kit 3.1 with 3’ Feature Barcode Kit and dual indexing (10x Genomics Protocol CG000316 Rev C or D). Libraries were sequenced as PE150 by Illumina sequencing to 65-80% saturation. Reads were mapped to the murine genome mm10 (GENCODE vM23/Ensembl 98) by using 10x Genomics’ cell ranger pipeline (6.0.0) using default settings.

### Quality control and normalisation of single cell RNA sequencing libraries

Two publicly available single cell RNA sequencing datasets of murine synovial stromal cells from STA models GSE129087 ^12^ and GSE145286 ^13^ well as the in-house datasets of synovial stromal cells were individually controlled to exclude low-quality cells using either the provided steps of the corresponding published study, or the criteria established for each individual dataset by investigating the distribution of unique molecular identifier (UMI) counts, number of detected genes, mitochondrial ratio, and the overall complexity (log10(genes/UMI)) (**Suppl. Tab. 1).** Using Seurat (4.1.1) ^14, 15^ and R (4.2.1), in-house datasets were additionally, prior to quality control, subjected to hashtag antibody oligos (HTO) separation using Seurat’s “HTODemux” with 0.99 threshold, in order to separate replicates or conditions, and to remove doublets and negative cells. Genes expressed in less than five percent of cells were discarded. Each dataset was individually processed by following the log normalisation workflow (“NormalizeData”, “FindVariableFeatures”, “ScaleData”, “RunPCA”) to test whether undesired sources of variations, such as cell cycle phase scores, tissue dissociation-induced stress scores ^16^ as well as mitochondrial ratio, were non-uniformly distributed in the PCA space, and therefore were regressed out in the next step. “SCTransform” (v2) ^17^ was then applied individually on the raw UMI counts of each sample.

### Integration of single cell RNA sequencing datasets

Five different integration strategies (Seurat CAA ^18^, Seurat RPCA ^14^, Harmony ^19^, Scanorama ^20^ and scVi ^21^) were applied and evaluated using k-nearest neighbour batch effect test (kBET) and batch-PC regression criteria ^22^. Harmony achieved the overall best performance in both criteria (**Suppl. Fig. 1a, b**). For Harmony integration, SCT transformed counts, along with the variable features and scaled data, were merged and subjected to PCA, before computing the harmony with top 50 PCs. Top 36 Harmony components, containing 90 percent of the total variation, were used for clustering and visualisation. The optimal resolution for clustering was determined using the co-dependency index-based (CDI) specificity measure (**Suppl. Fig. 1c**) ^23^. Using this resolution, 11 clusters were identified, including two endothelial cell clusters, pericytes, epithelial cells, five clusters of mesenchymal cells and the remaining *Cd45*+ cells (**Suppl. Fig. 1d, e**). Among the mesenchymal cells, *Pi16*+ cells and osteoblasts were identified using Seurat’s “FindSubCluster” method. Uniform manifold approximation and projection (UMAP) was created using the “RunUMAP” method. “PrepSCTFindMarkers” was then applied to the SCT assays to remove the variable sequencing depth effect, before running “FindAllMarkers” to identify markers of each cluster with default parameters. For downstream analyses hTNFtg197 + *I123*mc mesenchymal cells were subset from the integrated dataset and separately processed similarly but without the integration since UMAP visualisation did not show evidence of sample-specific clustering (**Suppl. Fig. 1f**).

### Analysis of R5+ ILC single cell RNA sequencing datasets

Single cell RNA sequencing datasets of synovial R5+ ILCs were subjected to HTO separation and quality control (**Suppl. Tab. 1**) as outlined above. A series of genes (**Suppl. Tab. 2**), biologically not relevant to our study, that were non-uniformly expressed and therefore influenced clustering, were removed from the dataset. Seurat’s “SCTransform” and CCA integration were used followed by PCA, UMAP, and clustering based on the integrated assay. Clusters of T cell subsets and *Cd45*-cells were excluded and the workflow of integration was repeated (**Suppl. Fig. 2a, b**). “FindClusters” with 0.6 resolution resulted in six clusters, one showing upregulation of precursors-associated biological processes, identified by GO enrichment analysis using Clusterprofiler (4.4.4) ^24^, as well as three ILC2s and two ILC3 subsets (**Suppl. Fig. 2c-e**).

### Single sample gene set enrichment analysis

A collection of gene sets (**Suppl. Tab. 3**) involved in arthritis damage, inflammation and resolution of inflammation selected from the Gene Ontology biological processes (GOBP), as well as Kyoto Encyclopedia of Genes and Genomes (KEGG), were used in the single sample GSEA (ssGSEA) analysis from Escape (1.6.0) ^25^, applied to the merged datasets of wildtype and hTNFtg197 + *I123*mc mesenchymal cells. Conserved differentially enriched pathways were identified using Wilcox rank sum test with Bonferroni correction on each condition, followed by averaging the log fold changes (LFC) over conditions, while taking the maximum adjusted p values as the overall significance criteria. Moreover terms with LFC ranging in the bottom 25 % quantile of the absolute LFC values were excluded. For visualisation, LFCs were scaled. For spatial transcriptomic data, Ucell scores of the genesets previously introduced in murine fibroblast subsets were calculated on the spatial and feature enhanced spatial dataset, and were subjected to Wilcoxon rank sum test between the spatial clusters. Resulting LFCs were scaled, and terms with Bonnferoni adjusted p-value < 0.05 were selected for visualisation.

### PseudoBulk differential gene expression

To perform differential gene expression (DGE) tests between conditions, pseudo bulk counts matrices were created by extracting and aggregating the UMI counts over replicates and cell types, and DGE was performed using the DESeq2 wald test ^26^.

### Receptor internalisation

DGE was performed between untreated and anti-mIL-17a conditions in the murine hTNFtg197 + *I1*23mc synovial mesenchymal cell dataset. LFC sorted results were subjected to fGSEA (1.22.0) ^27^ using 71 subterms of endocytosis from GOBP (GO:0006897). The top five terms (negatively or positively enriched) were selected for visualisation, significance was highlighted when the p-values were smaller than 0.05. To study correlation of receptor internalisation with damage and resolution activity, its UCell ^28^ score was computed by taking the leading edge genes of the “positive regulation of receptor internalisation” term from the fGSEA result. Damage and resolution UCell scores were assigned by taking the genes intersecting between damage- or resolution-associated pathways (**Fig. 4a**), and the top markers (adjusted p-value < 0.05 and LFC > 1) of the respective fibroblast subsets. These scores were then averaged over replicates in each fibroblasts subset, and subjected to Pearson correlation with Benjamini-Hochberg correction.

### RNA velocity estimation and directional fate mapping

Velocyto workflow (0.17.17 on Python 3.10.2) ^29^ for 10x Chromium generated single cell RNA sequencing was applied to the sorted binary alignment files, to create the unspliced and spliced RNA count matrices for mesenchymal and ILC datasets. The scVelo (0.2.4 on python 3.9.10) ^30^ pipeline was implemented in the dynamic mode to estimate RNA velocities and visualised on the UMAP by using the method “velocity_embedding_stream”. To further quantify the differentiation probabilities, using CellRank’s (1.5.1 on python 3.9.10) ^31^ Generalised Perron Cluster Cluster Analysis (GPCCA) estimators, an aggregated transition probability matrix was generated by combining velocity and transcriptional similarity kernels (equally weighted). After assigning terminal states by “compute_macrostates”, using “compute_absorption_probabilities”, the probability of differentiation of each single cell was calculated, and visualised using “plot_absorption_probabilities”. These probabilities were averaged for each condition and toward each fate individually in each of the sublining fibroblast subsets and visualised in a heatmap.

### Differential abundance analysis

edgeR (3.38.1) ^32^ was used to perform the cluster differential abundance test. Cluster counts in all conditions and in each replicate were prepared as input to edgeR followed by “estimateDisp”, “glmQLFit” and “glmQLFTest”^33^. In each test those with a false discovery rate < 0.05 were considered significant.

### Sample-associated relative likelihood (MELD)

MELD (1.0.0 on Python 3.8.8) ^34^ was used to quantify treatment effects on the cellular composition. PHATE dimension reduction was performed followed by PCA. Replicate associated densities were calculated by running “meld”with default parameters. These densities were normalised (L1 normalisation to enforce cell wise sum to be 1) and averaged over all replicates in each condition per cell, and their values were visualised as condition-associated relative likelihood in major clusters of *Cd200*+ vs *Cd200*-, as well as in all FAP expressing subclusters.

### Gene co-expression network analysis

To identify and analyse correlated gene programs in the single cell RNA sequencing dataset of ILCs, hdWGCNA (0.2.02) ^35^ was used by grouping transcriptionally similar cells separately in each replicate and each subcluster of ILCs. “TestSoftPower” was applied to identify the value to which the correlations were raised, and highly correlated modules were identified using the “PlotDendrogram” function. To visualise the co-expression network, we calculated the UMAP embedding of the topological overlap map using “RunModuleUMAP” and “ModuleUMAPPlot” methods.

### Receptor-ligand interaction analysis

CellChat R toolkit (version 2.2, R 4.0.1) ^36^ was used to explore cellular interactions between mesenchymal cell subsets and ILCs. Communication probabilities were computed using the cellChats datasets of secreted and cell-cell contact signalling. Condition-specific enriched interactions were identified by merging the resulting CellChat objects, “rankNet” function followed by row-normalisation was applied, and pathways with more than 80 % relative contribution were selected (**Suppl. Fig. 2f**). The enriched interaction network was extracted from each of the CellChat objects, and was visualised using chord diagrams, with thickness of the arcs corresponding to the total interaction strength.

### *In silico* perturbation analysis

To simulate genetic knock-out of *Cd200r1* and to study its effect on ILC differentiation, Dynamo’s (1.1.0, Python 3.9.10) *in silico* perturbation method was implemented ^37^. The integrated dataset of ILCs was subject to the standard preprocessing step (“recipe_monocle”), followed by recovering velocity dynamics, and reconstructing the vector field on the basis of PCA. “Dynamo.pd.perturbation” was used by setting the *Cd200r1* expression to −100 and the perturbed vector field was visualised using the “streamline_plot”.

### Human PsA single cell RNA sequencing analysis

Two publicly available datasets of human PsA single cell RNA sequencing from whole synovial tissue ^38^ and CD45+ sorted cells ^39^, were analysed. Quality control included exclusion of low quality cells with the criteria UMI count < 1000-500, number of features < 500-250, mitochondrial ratio > 0.25-0.12 and complexity < 0.95 - >0.8, and by removing genes expressed in less than five percent of the cells as well as mitochondrial, ribosomal and stress associated genes. Seurat’s log normalisation workflow was performed while regressing out cell cycle scores and UMI count. After PCA calculation, the top 20 PCs were used for Harmony integration. Top 20 harmony components were used for clustering and visualisation.

### Integration of mouse mesenchymal and human PsA fibroblast

To identify the human counterpart of the murine fibroblast subtypes both datasets were integrated. Human PsA fibroblasts were subsetted and genes translated into their murine orthologs, merged with mesenchymal cells from the murine hTNFtg197 + *I123*mc model. After Seurat’s “SCTransform” of each sample individually, CCA integration was performed to overcome the strong interspecific bias, followed by PCA, clustering and UMAP visualisation. Under the assumption that cells with higher transcriptional similarity colocalize in the same cluster, human counterpart of murine *I16*+, *Mmp3*+ and *Cd200*+ fibroblast subsets were identified by plotting the percentage of each subset in the identified integrated clusters.

### Generation and sequencing of spatial transcriptomic libraries

Unfixed human synovial biopsies were snap frozen in OCT Tissue TEK (Sacura). 10 μm thin sections were cut on a Cryotome FSE (Thermo Fisher) to generate spatial transcriptomes according to the Visium Spatial Gene Expression Kit (v1 chemistry, 10x Genomics). An optimization slide was run for optimal permeabilization conditions. 6 min was found to be optimal. Spatial transcriptomes were prepared according to the Visisum Protocol CG000239 Rev B: Libraries were sequenced as PE150 by Illumina sequencing to >80% saturation.

### Preprocessing and spatial clustering

Reads were mapped to the human genome GRCh38 (GENCODE v32/Ensembl 98) by using 10x Genomics’ space ranger pipeline (1.2.1) with default settings manual correction of spots under tissue using Loupe Cell Browser (5.0.1). BayesSpace (1.0.1) ^40^ workflow was applied to analyse spatial transcriptomic samples. “readVisium” was used to import count and spatial matrices followed by an overall filtering of regions of the tissue with very few cells, and spots with < 500 genes and < 4,000 UMI counts, as well as with > 25 percent mitochondrial ratio. “spatialPreprocess”with default parameters was applied. “qTune” and plotting suggested an elbow of eight for optimal number of spatial clusters (**Suppl. Fig. 3a**). “spatialEnhance” and “enhanceFeatures” with top 2,000 variable genes from the single cell RNA sequencing dataset as well as the selected list of markers, was used to enhance the spot and feature resolutions. Spatial cluster 3 with relatively lower UMI count, included regions with little tissue or border regions (**Suppl. Fig. 3b, c**), and was discarded.

### Gene module scores and hierarchical clustering

UCell scores were computed for each of the cell-type gene modules, identified either in the single cell RNA sequencing analysis of PsA or from the literature (**Suppl. Tab. 4**). These scores are then averaged and scaled over all different single spots using Seurat’s “DotPlot”, grouped by the spatial clusters, and were subjected to row-wise and column-wise hierarchical clustering using Pheatmap ^41^.

### Imaging mass cytometry

Human synovial biopsies were fixed in PBS containing 4 % (w/v) formaldehyde for 16 - 24 h. Tissue was dehydrated and infiltrated with paraffin, embedded in paraffin blocks, cut to 5 μm thin sections and mounted on slides. Paraffin embedded thin sections were deparaffinized by heating the slides for 30 min at 65 °C and washing in Histo Clear (National Diagnostics) and rehydrated with a series of 100 % (v/v) ethanol, 95 % (v/v) ethanol, 80 % (v/v) ethanol, 60 % (v/v) ethanol, water. Heat induced epitope retrieval was performed for 30 min at 95°C in Tris-EDTA buffer (10 mM Tris, 1 mM EDTA, pH 9.2). After cooling down for 15 min, blocking was performed in TBS-T supplemented with 3 % (w/v) BSA and 1% rabbit serum for 1 h at ambient temperature. A precomposed panel of metal labelled antibodies (**Suppl. Tab. 5**) was stored at −80°C and applied to the sections after thawing and centrifugation at 15,000 g for 10 min. Incubation with antibodies occurred over for 16 h at 4°C in presence of 0.3 μM Ir-intercalator (Standard Biotools). Slides were washed twice in TBS-T after staining for 10min, followed by a brief wash in deionised water, air dried and kept in a desiccated environment until analysis. Acquisition was run at 200 Hz on a Hyperion Imaging System (Standard Biotools). MCD files were processed according to the IMC segmentation pipeline (3.5) ^42^. Spectre ^43^ was used to convert TIFF files into FCS files, which were then analysed with FlowJo (v10.8.1, BD Biosciences).

### Statistical analysis

Statistical analysis on non RNA sequencing data was performed as described in each section using Prism 8 software. Unless otherwise stated, all data are presented as mean ± s.d. and obtained from at least two independent experiments. Parametric and non-parametric analyses were used where appropriate based on testing for a normal distribution using the D’Agostino-Pearson omnibus normality test. Differences were considered to be significant when *P* <0.05. Multiple testing corrections were applied where appropriate.

## Data availability

Single cell sequencing data that support the findings of this study have been deposited in Gene Expression Omnibus (GEO) with the accession codes GSEXXXXXXX. Source Data for are provided with the online version of the paper.

## Code availability

All the methods and algorithm used in this manuscript are from previously published studies and cited in the methods section. Selection criteria, threshold values, and other essential parameter are stated in the methods section. No new method was developed to analyse data. Additional scripts to reproduce the analyses are available from the authors upon request.

## Supplemental Figure Legends

**Supplemental Figure 1 (a)** Scatter plot showing the integration assessment scores from five evaluated integration strategies (scVI, Seurat RPCA, Scanorama, Seurat CCA, Harmony) for the cross-study stromal single cell RNA sequencing. **(b)** U-map visualizing the distribution of cells of each of the harmony integrated datasets. **(c)** CDI specificity measure for different resolutions of Seurat’s clustering algorithm. **(d)** U-map of the integrated dataset visualizing clustering at resolution 0.2 and annotation of corresponding cell types. **(e)** Heat map of the top ten genes expressed in the 11 identified clusters of the integrated dataset. **(f)** U-map visualization of the synovial mesenchymal cells from the hTNFtg + *I123*mc datasets without integration.

**Supplemental Figure 2 (a)** U-map visualizing the major cell types identified in the single cell datasets of R5+ sorted lineage negative cells (ILCs). **(b)** T cell marker genes visualized on the U-map. **(c)** U-map visualizing the identified subclusters of ILCs after removal of contaminant cells. **(d)** GOBP functional enrichment analysis on the significantly upregulated genes in the ILC precursors. **(e)** Visualization of selected markers of type 2 and type 3 ILCs on the U-map. **(f)** Significant interaction pathways identified with CellChat in the merged ILC-Fibroblasts single-cell dataset of STA and STA + *I123*mc models. Labels are coloured with respect to the condition when the relative contribution of the interaction pathways is > 80 %.

**Supplemental Figure 3 (a)** BayesSpace average negative log likelihood for different numbers of spatial clusters on the Visium spatial transcriptomics dataset of PsA. **(b)** Violin plot showing the distribution of UMI counts for each spatial clusters. **(c)** Distribution of spatial clusters overlayed on the Visium H & E stained slide.

