## Supplementary figures and images for "Molecular Imaging with Fibroblast Activation Protein Tracers depicts Inflammatory Joint Damage and its Transition to Resolution of Inflammation"

### Suppl. Fig. 1

#### Supplemental Figure 1

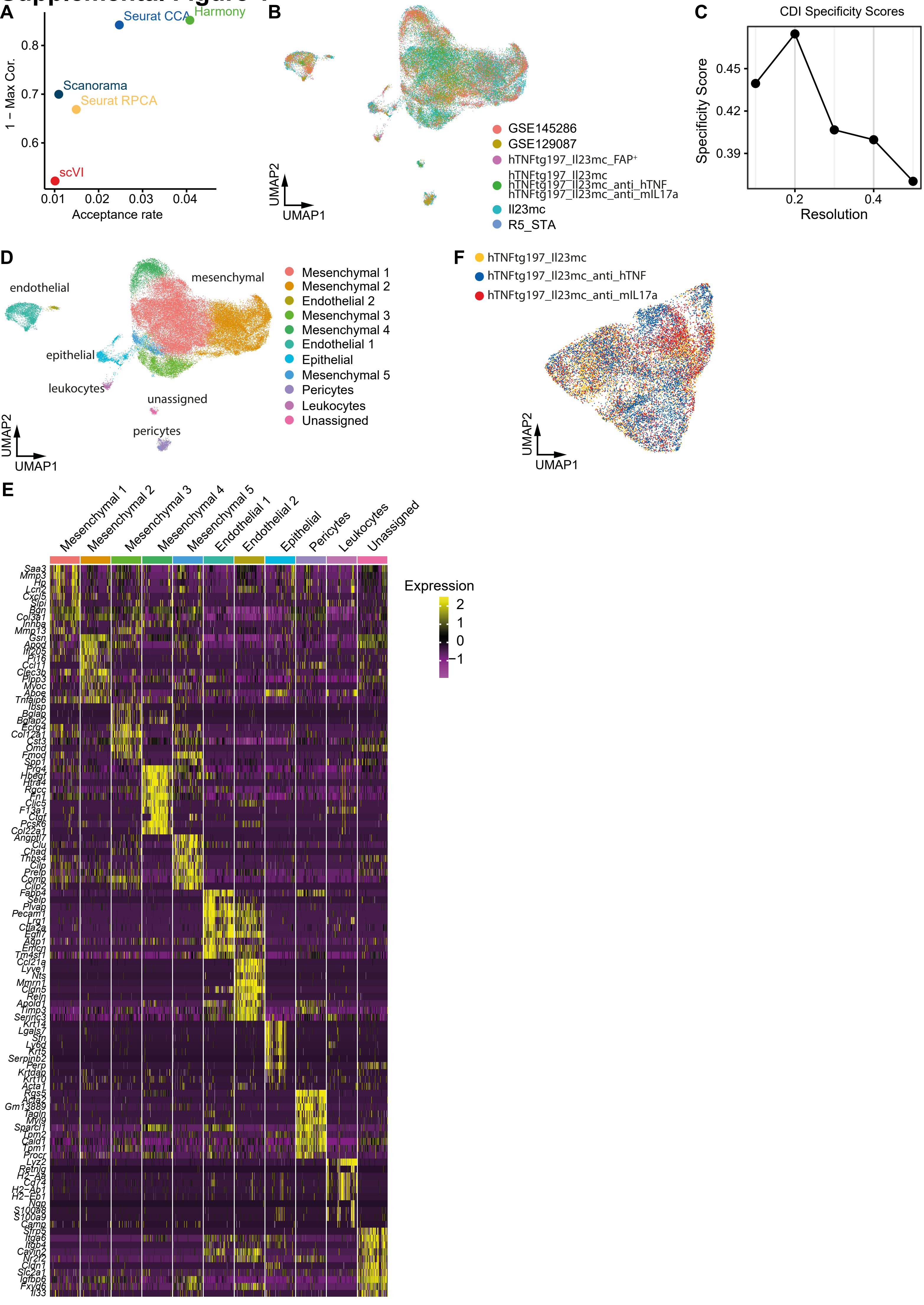

### Suppl. Fig. 2

#### Supplemental Figure 2

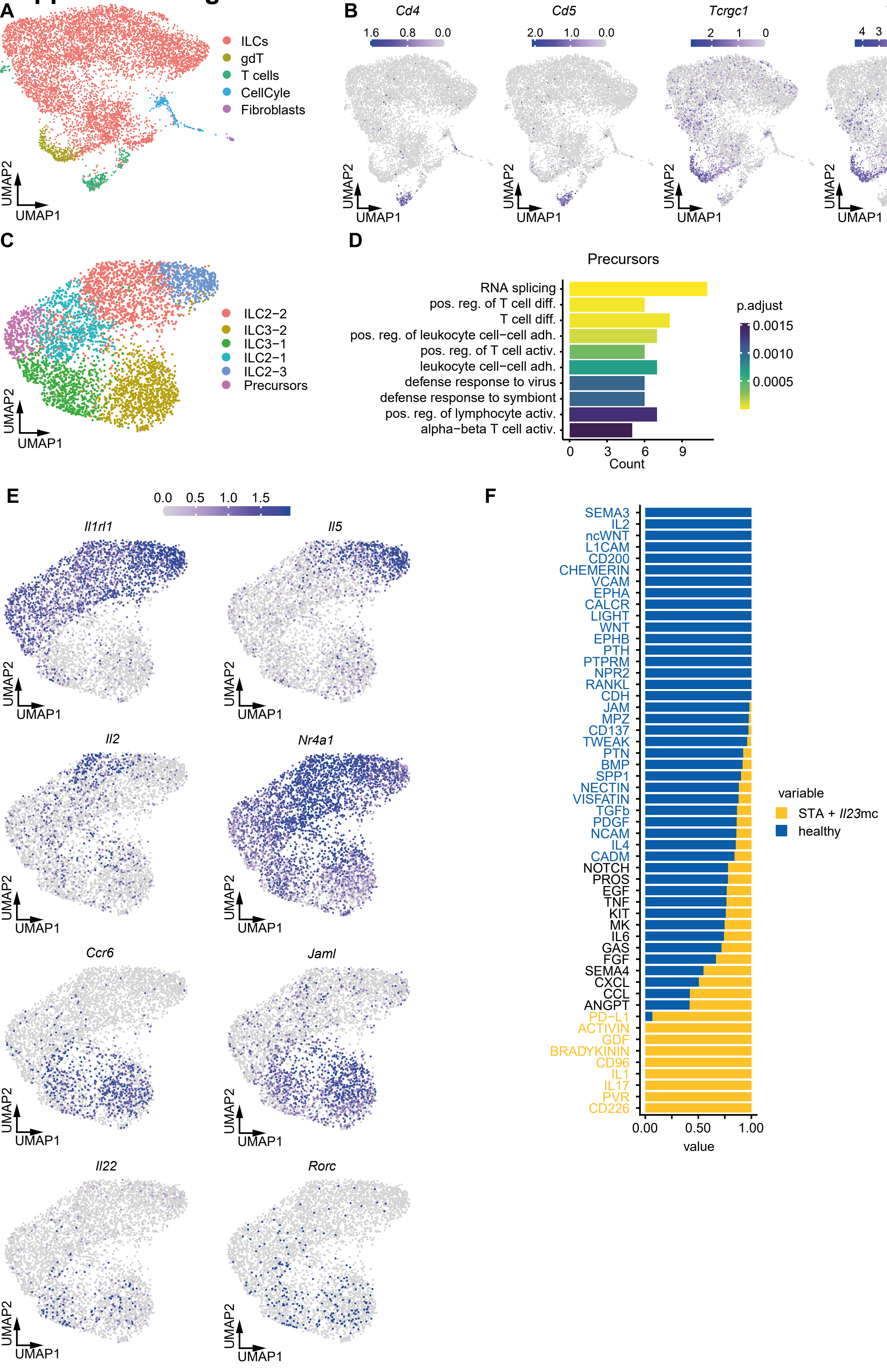

### Suppl. Fig. 3

#### Supplemental Figure 3

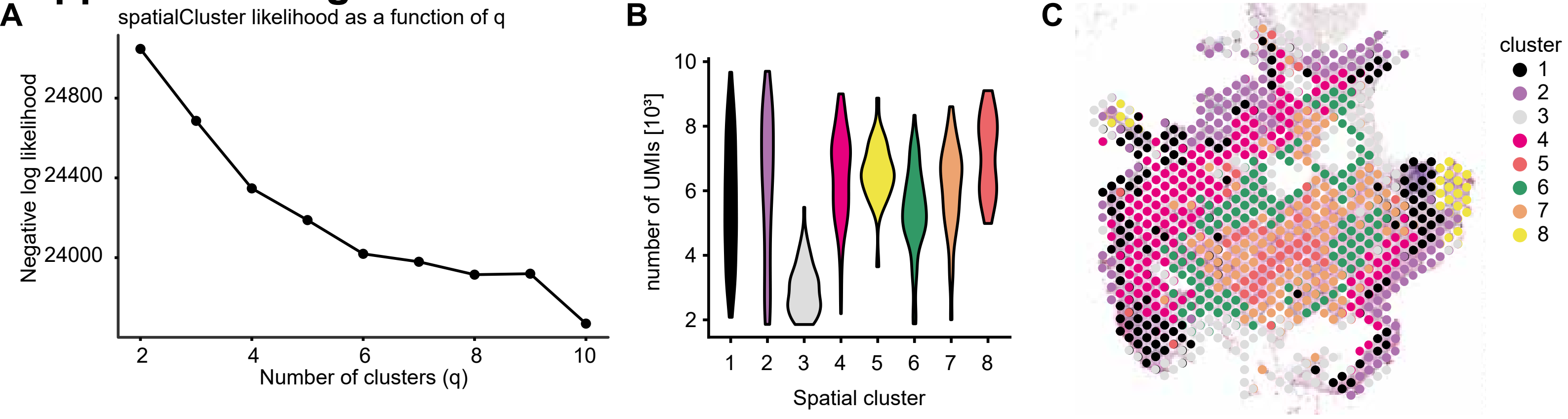
